# RNA Targets and Physiological Role of Topoisomerase 3β

**DOI:** 10.1101/2023.05.25.542246

**Authors:** Shohreh Teimuri, Beat Suter

## Abstract

Topoisomerase 3β (Top3β) works not only on DNA but also RNA. We isolated and identified the naturally cross-linked RNA targets of Drosophila Top3β from an early embryonic stage that contains almost exclusively maternal mRNAs. Favorite targets were longer RNAs, RNAs with many splice sites, and RNAs that become localized in large cells. Demonstrating the importance of its enzymatic activity, Top3β without the hydroxyl group that makes the covalent bond to the RNA, did not allow normal expression and subcellular localization of tested gene products of the identified targets. *Top3β* is not essential for development to the adult stage but to maintain the morphology of the neuromuscular junction in adult flies and to prevent premature loss of coordinated movement and aging. Alterations in human *Top3β* have been associated with several neurological diseases and cancers. The homologs of genes and (pre)mRNAs mis-expressed in these conditions show the same characteristics identified in the Drosophila *Top3β* targets, suggesting that Drosophila could model the function of human *Top3β*. Indeed, an *in vivo* test of this model showed that the enzymatic activity of Top3β reduces the neurodegeneration caused by the human (G4C2)_49_ RNA. Together, these studies show that *Top3β* plays important roles in normal gene expression, particularly for large genes and long and complex transcripts that need to be transported and translationally controlled. Its absence stresses the cells, seemingly increasing the chances of contracting various neuronal and neurodegenerative diseases and cancers.

## Introduction

Topoisomerase 3β (Top3β) is an RNA-binding protein (RBP) with dual enzymatic activity toward DNA and RNA (1) (88). Topoisomerases are better known for their activity toward DNA, where they resolve tension and supercoiling, a prerequisite for efficient transcription, transcription control, and replication (11) (86) (recently also reviewed in 87). Much less is known about the function of topoisomerases toward RNA. Along with other RBPs, Top3β interacts with the RNAi machinery (49), and it contributes to mRNA stability and translation, particularly in the nervous system, which is significant for neurodevelopment and psychological and physical well-being (2)(74)(85). Its functional importance is suggested by results that link impaired *Top3β* function to various neurodevelopmental and cognitive disorders (2)(3)(73)(84).

*Drosophila melanogaster* Top3β possesses the key features of a topoisomerase. The predicted catalytic tyrosine (Tyr, Y; (4) (90)) residue is present at position 332 (Y332). Top3β topoisomerases distinguish themselves from other topoisomerases by their conserved RNA-binding motif called RGG (Arginine Glycine Glycine) box, which binds to RNAs or proteins. Its paralogue Top3α, a DNA topoisomerase, possesses the catalytic tyrosine residue but lacks the RGG box (4). *In vitro*, the RGG box is not required for the enzymatic activity but might play a role in binding to the RNA and thereby facilitating the enzymatic activity *in vivo* (2).

We hypothesized that the neurological requirements for an RNA topoisomerase may in part reflect its activity on mRNAs that need to be transported over long distances to be translated at the right place and time. During such transport, long molecules, such as mRNAs, are likely to become entangled or suffer other structural problems that might interfere with their packaging, transport, unpackaging, and translation (1)(94). Therefore, a topoisomerase that acts on RNAs might be able to rescue such RNAs. As discussed by Su and colleagues (74), there is considerable evidence that Top3β activity on RNA has important functions in cells and that it might act at all levels of gene expression. Using human cell lines, these authors showed that Top3β indeed affects mRNA translation and RNA stability (74). They also found evidence that these effects involve mainly TDRD3, which could be strengthened by the recently described mouse *Tdrd3*-null phenotype (85). However, there is still surprisingly little knowledge about the steps in gene expression where Top3β acts on RNAs and particularly about the *in vivo* role of this activity for the cells and the organism. Especially in large cells, where mRNAs need to travel over long distances, an RNA topoisomerase could provide an important advantage and allow the cell to function properly. Drosophila has such large cells in the nervous system, the female germline, and the young, syncytial embryo. The latter is easily accessible and free of additional tissue. According to Flybase (https://flybase.org/reports/FBgn0026015), the expression of Drosophila Top3β is particularly abundant in 0-2 hours old embryos, when the syncytial embryos are in the first 10 nuclear divisions cycles (78). The mRNAs in these large cells are maternally provided and bulk zygotic transcription would only start about an hour later, around cellularizations (reviewed by (79), (80)).

Proper mRNA localization to specific compartments combined with controlled local translation is essential for normal differentiation and cellular physiology and it has been studied extensively during these stages. This developmental stage, therefore, appears to provide a unique opportunity to study the activity and function of Drosophila Top3β toward cytoplasmic RNAs by identifying relevant *in vivo* RNA targets.

Here we show that in such 0-2 hours old embryos, longer RNAs were more frequently bound to Top3β in a Y332-dependent way. This is consistent with the notion that larger RNAs are more likely to encounter structural problems that are being targeted and repaired by the RNA topoisomerase in living cells. The same RNA features are also over-represented among the differentially expressed RNAs in the *Top3β* mutants. As opposed to the capturing of the targets in young, preblastoderm embryos, the transcriptomics analysis is expected to reveal the sum of direct and indirect effects of the cytoplasmic and the nuclear activities of Top3β. Specific mutations of the Y332 active site tyrosine (Y332F) and the RGG box (ΔRGG) revealed the importance of the enzymatic and RNA/protein binding activity, respectively. The enzymatic activity of Top3β is needed to covalently bind to mRNAs and for the normal accumulation of many long mRNAs and their encoded proteins in general, and also in specific subcellular locations. The lack of this enzymatic activity is not essential for the viability of the embryos or larvae but stresses the affected cells and causes neurodegeneration and premature aging of adults. Because the Drosophila enzyme affects the outcome of a human RNA that causes neurodegeneration, this Drosophila model produces valuable novel insights into neurodegenerative diseases and cancers with which human *Top3β* had been associated.

## Results

### RNAs depending on *Top3β* for their normal expression

To test whether Top3β affects gene expression at the RNA level, RNAs from 0-2 hours-old embryos were extracted from a wild-type control strain and the *Top3β* mutants *Top3β^26^*, a null mutant (49)(50) caused by a deletion of a large fraction of the coding sequence starting at codon Y141 and changing it into a stop codon, *Top3β^Y332F^*, and *Top3β^ΔRGG^*, point mutants generated for this work to test the function of Y332 and the RGG motives. All mutants were both maternally and zygotically mutant. Sequence analysis of the transcriptome carried out on biological triplicates established the effect of the different *Top3β* mutations on the RNA levels expressed in 0-2 hours old embryos (**Supplementary Table S1-3; Supplementary Fig. S1**). Using adjusted p-values (Adjp) <0.05 and log2 fold changes <-1 and >1, RNAs that showed level changes between the mutant and the wild-type strains were selected for a first analysis (**Fig. 1A**, left and central panel). Much more stringent conditions (Adjp <0.0002 and log2 fold changes <-1) were also used (**Fig. 1A**, right panel). In both analyses, the strongest effect was observed with the null mutant *Top3β^26^*which also displayed a very strong reduction of the *Top3β^26^* RNA levels (**Supplementary Table S1; Supplementary Fig. S1).** As opposed to the null mutant, the RNA with the point mutation was stable **(Supplementary Tables S2-3)**. The gene ontology analysis of biological processes revealed similar results for *Top3β^Y332F^* and *Top3β^26^*, but no enrichment terms for *Top3β^ΔRGG^*(**Supplementary Fig. S2).** The affected processes range mainly from morphogenesis, synapse signaling, and trans-membrane transport to cell adhesion and cuticle formation. Several of these processes are active repeatedly in the life cycle, and this includes neurons and their synapses.

**Fig. 1.**
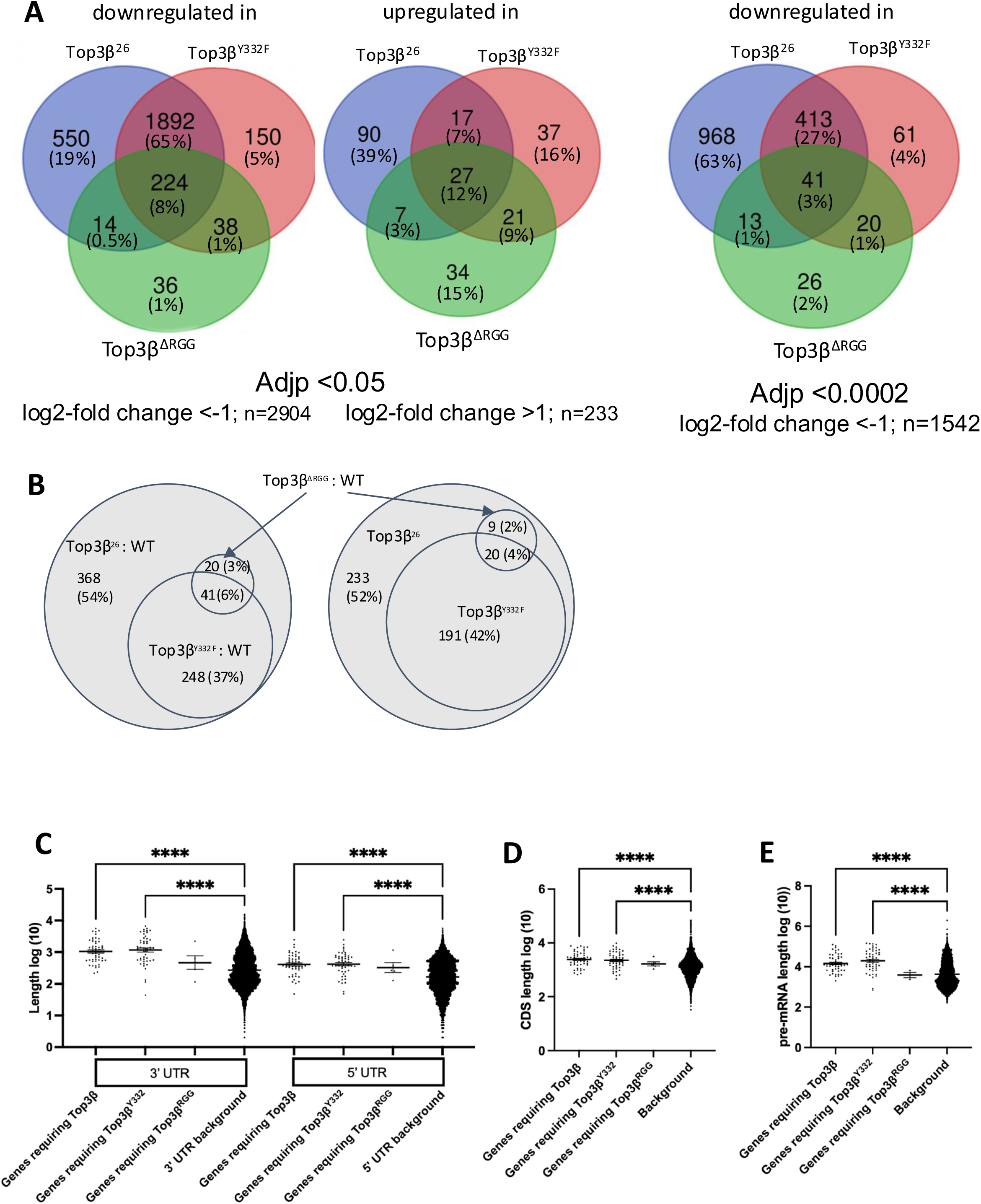
Effect of the *Top3β* mutations on the transcriptome of 0-2 hrs. old embryos. **A)** The left and central panels show the changes compared to the wild-type (WT) transcriptome using adjusted p-values (Adjp) <0.05 and log2-fold change <-1 (downregulated) and >1 (upregulated). The Venn diagram shows also the allocation of the data to the 3 different mutants. Using the more stringent criteria Adjp <0.0002 and log2-fold change <-1, the right Venn diagram shows the fractions of the genes with reduced transcript levels and which mutant showed the reduction. n: is the number of different genes identified in the three mutants. **B)** Differentially expressed transcript isoforms in the *Top3β^26^* null mutant compared to the wild type (WT). Adjp <0.05 and absolute log2-fold change ≥1. The Venn diagram shows which fraction is also differentially expressed in the point mutants when analyzed with the same stringency parameters (left panel). The fractions of the differentially expressed transcript isoforms expressed at lower levels in the mutants are shown in the right diagram. **C)-E)** RNA features of the genes, which produce the transcript isoforms, that depend on *Top3β* to reach normal expression levels. **C)** Transcript isoforms from genes that produce long UTRs are over- represented in the group of RNAs that depend on *Top3β* and its Tyr^332^ (Y332) residue to reach normal expression levels. **D)** Transcript isoforms from genes that encode longer CDS are expressed at higher levels in the wild type than in the *Top3β^26^* and *Top3β^Y332F^*mutants. **E)** Transcript isoforms from genes that produce longer pre-mRNAs are expressed at higher levels in the wild type than in the *Top3β^26^*and *Top3β^Y332F^*. Expression levels were compared to the *Top3β^+^*(WT) control in 0-2 hrs. old embryos. p-value <0.0001=****, p-value <0.001=***, p- value <0.01=**, p-value <0.05=*. n=50 for the null mutant *Top3β^26^* and *Top3β^Y332F^* and n=5 for *Top3β^ΔRGG^*.

Only a small fraction of the affected genes displayed upregulation in the mutants (**Fig. 1A; Supplementary Fig. S3**). The majority were downregulated. The downregulated genes still showed a high overlap between the point mutations and the null mutant, indicating that the point mutations act as (partial) loss-of-function mutations. 92% and 85%, respectively, of the effects seen with the Y332F mutant using the Adjp <0.05 and Adjp <0.0002, respectively, were also observed with the null mutant. This points to the importance of the active site Tyrosine (Y332) for the function of Top3β and indicates that the enzymatic activity is important for the proper expression levels of many genes. The strong overlap with the null mutant is also reassuring because the two mutants are from different genetic backgrounds. In contrast, the *Top3β^ΔRGG^* mutation was induced in the same genetic background as the *Top3β^Y332F^* mutation. The fact that with both thresholds only 11% of the *Top3β^Y332F^* hits were also identified in the *Top3β^ΔRGG^* mutant shows that strain background differences do not, or only weakly, affect the data quality.

Large genes tend to produce more frequently different mRNA isoforms, sometimes with different protein-coding capacities and even functions. With the standard expression profiling used so far, we might miss a strong effect on one isoform if other mRNA isoforms are not affected or compensate for the missing one. To find out whether *Top3β* is needed to produce normal levels of specific mRNA isoforms, we allocated the sequence hits to specific transcripts using the Salmon method (91) (**Supplementary Tables S4-S7, Fig. 1B**). The expression levels of individual transcript isoforms were then compared between the null mutant and the wild type using as cut off Adjp <0.05 and absolute log2 fold changes ≥1. This yielded 677 differentially expressed RNAs of which 67% (453) were downregulated (and 224 upregulated) in the null *Top3β^26^* mutant, suggesting again that *Top3β* is mainly needed for the production or stability of RNAs. The transcript isoforms that are upregulated in the mutant, might either be direct targets or generated by indirect, e.g., compensatory effects. A closer inspection revealed that many genes with a transcript species that was upregulated in the mutant had a different isoform that was downregulated in the same mutant. This is consistent with a compensatory mechanism and a genomic background effect, but also with the lack of Top3β affecting splicing, polyadenylation site selection, or promoter usage, causing the expression to shift to a different RNA isoform.

We next investigated the contribution of the catalytic Tyr, which is essential for the enzymatic activity, and the RGG motives, which mediate interactions with RNAs and proteins, toward the expression of specific transcript isoforms. To reduce strain-specific and random effects, we used the same cut-off values as before and focused on the 677 genes that are differentially expressed in the null mutant. As shown in **Fig. 1B**, close to half (43%) of the transcript isoforms identified in the null mutant were also identified as differentially expressed in the *Top3β^Y332F^* mutant (**Supplementary Table S7**). In the *Top3β^Y332F^* mutant, 211 out of 289 (73%) differentially expressed RNAs were downregulated in the embryos, suggesting that the enzymatic function acts mainly to support RNA expression. Furthermore, this fraction is close to the 67% observed in the null mutant, indicating that the catalytic mutant has similar effects on the transcriptome of 0-2 hours old embryos as the null mutant. The impact of the mutated RGG domain on the embryonic transcriptome was less pronounced, with 29 out of 61 (47%) RNAs downregulated in the embryo, many might be indirect targets. Remarkably, 20 of these 29 RNA isoforms depended not only on the RGG motives of Top3β but also on the Y332 for their normal expression levels **(Fig. 1B)**. Changes in mRNA levels in 0-2 hours old embryos of *Top3β* mutants are not only due to a combination of direct and indirect effects of a specific mutation but also reflect the combined effects of the Top3β activity on the DNA of the nurse cells during oogenesis and on the pre-mRNAs and the mRNAs during oogenesis and embryogenesis.

We next selected from the list of preselected transcript isoforms **(Supplementary Table S7)** the 50 that showed the strongest reduction (according to their log2fc) in the null mutant and the Y332F mutant, and the top 5 for ΔRGG that were also in the top list of the null mutant. We then asked which RNA features were overrepresented in this group compared to the wild-type control. Selecting the top hits should reveal with higher confidence gene- and RNA features requiring *Top3β* activity to reach normal expression levels. On the other hand, many RNAs that appeared interesting according to the volcano blots were not included anymore (**Supplementary Fig. S1**). In embryos, the RNAs that were less expressed in the *Top3β^Y332F^* and *Top3β^26^* mutants had longer untranslated regions (UTRs), protein-coding sequences (CDS), and pre-mRNAs (**Fig. 1C-1E**). We conclude that long pre-mRNAs and mRNAs depend more strongly on *Top3β* to reach their normal expression levels.

The analysis of the significant changes in transcript isoform levels (**Supplementary Table S7)** revealed that at least 49 genes with reduced RNA levels of one isoform had at least one other isoform that was significantly higher expressed. Because these pairs might help to reveal the mode of action of *Top3β,* we studied 20 reciprocally expressed isoform pairs (**Supplementary Table S14**). In 11 cases, the upregulated transcript isoforms started more upstream, 3 further downstream, and 6 at the same location. The latter group displayed alternative and additional splicing and 3’end formation. Top3β is a good candidate for modulating such splicing activities. Due to the different start locations observed in 14 cases, the pairs with the different transcription start sites differed in their 5’UTR and sometimes in the N-term of the ORF, too. Additionally, differences in splicing and splice site selection were also observed. Some pairs displayed different protein-coding capacities while others did not. The *Shaker cognate l-RA* (*ShaI-RA*) isoform was present at higher levels in the *Top3β* null and Y332F mutants, but contains a shorter last intron, causing earlier translation termination and lack of C-terminal parts of the voltage-gated K-channel. *Shal* is involved in neuronal physiology, locomotion, and lifespan. Another example of strong differences between the pairs of mRNA isoforms was found in *Tropomyosin-1 (TM1*). Apart from 3’UTRs of different lengths, excision of a part of the 5’UTR, and different inclusions of small exons, the transcripts *TM1-RM* and *TM1-RA* start far apart and show very different alternative splicing patterns such that a large fraction of their ORF stems from different parts of the gene. Even though topoisomerases are involved in remodeling chromatin, the upregulation of transcript isoforms that initiate at different sites could also be the product of more indirect effects that compensate for the lack of another isoform.

### *Top3β* in neurological diseases and cancers

*Top3β* has been associated with several neurological diseases and cancers in humans. To test whether the human *Top3β* might similarly act to support the expression of higher levels, particularly of long RNAs, a list of diseases associated with *Top3β* was retrieved using gene-disease associations (DisGeNET (v7.0)(5)). The 12 diseases identified were either neurological diseases or cancers. The lists of differentially expressed genes associated with each of these diseases were then obtained from the same database and used to identify their *Drosophila* orthologues from DIOPT Version 9.0 (**Supplementary Table S8**). The list of these *Drosophila* genes was then compared with the list of transcript isoforms (and their genes) whose embryonic mRNA levels were reduced in the absence of *Top3β* (**Supplementary Table S7**). A sizable fraction of the 453 genes encoding *Top3β*-dependent isoforms, were also Drosophila orthologs of genes affected by these neurological diseases and cancers (**Fig. 2**). Due to the small number of genes affected in the Juvenile Myoclonic Epilepsy (JME), this disease was not considered for further studies. For the remaining ones, there is a clear overlap between the two datasets and the overlap ranges between 2 and >3 times the calculated random overlap, suggesting that the effects seen in the fly model with the inactivated *Top3β* also contribute to the human disease phenotype.

**Fig. 2.**
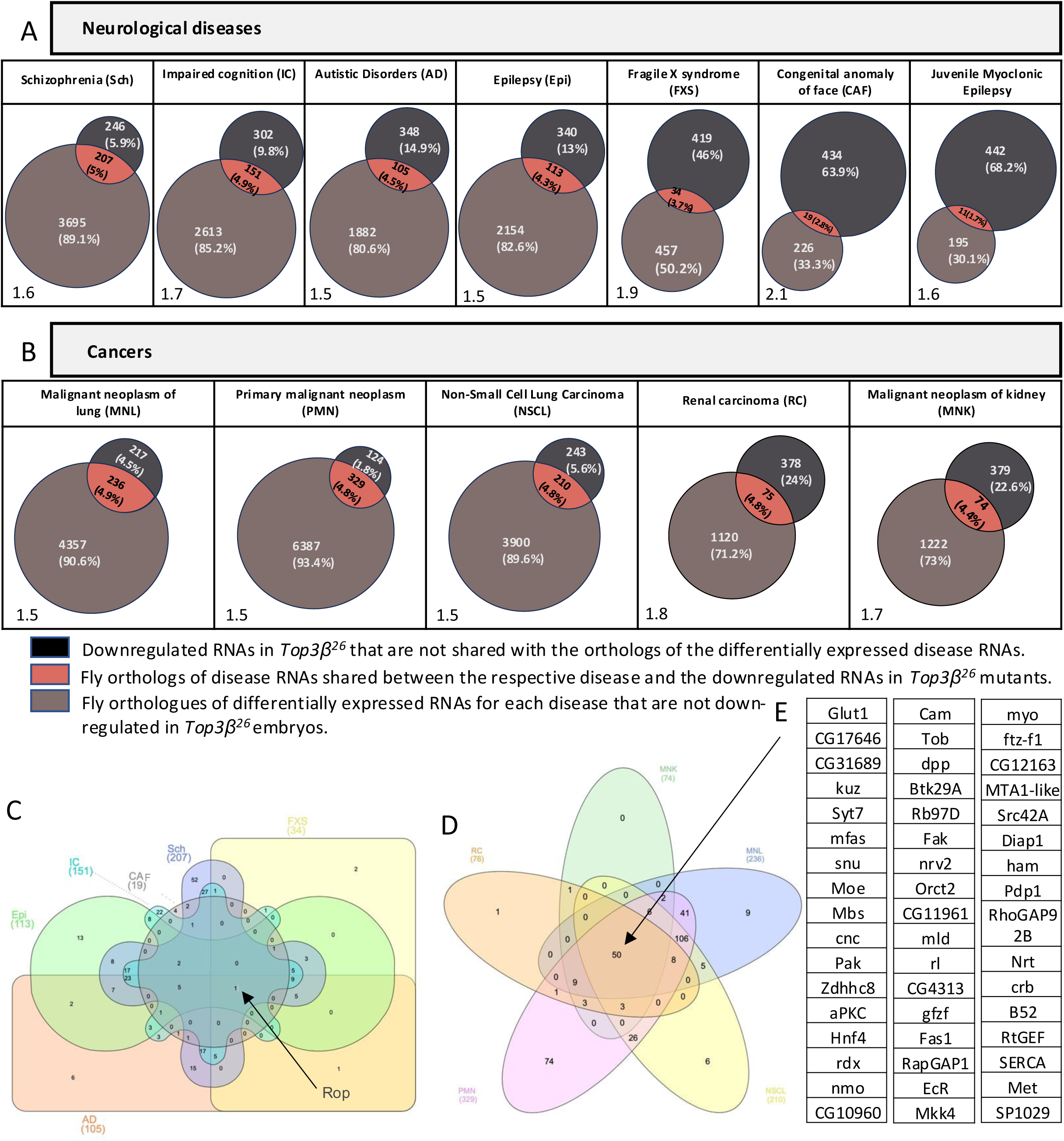
Overlap between downregulated embryonic RNAs in *Top3β^26^* and the fly homologs of differentially expressed RNAs in several cancers and neurological diseases associated with human *Top3β*. A, B) Venn diagram of the genes shared between the list of down-regulated RNAs in *Top3β^26^*embryos and the fly homologs of differentially expressed RNAs of the listed diseases. Numbers and frequencies are listed. The probability of random overlap is shown in % at the bottom left of each Venn diagram. **C)** Drosophila genes depending on *Top3β* for normal expression levels in embryos were placed into a Venn diagram such that their position shows in which neurological disease their homologs differentially expressed (note that JME was not included). *Rop* appeared in all lists. **D)** Drosophila genes depending on *Top3β* for normal expression levels in embryos were placed into a Venn diagram such that their position shows in which cancer their homologs are differentially expressed. **E)** List of the 50 genes that depend on *Drosophila Top3β* for their normal expression levels in embryos and whose homologs are also affected in all listed cancers associated with human *Top3β* alterations.

Notably, a single gene, *Rop* (*Ras opposite*), was present in the lists from all neurological diseases and the downregulated embryonic RNA isoforms (**Fig. 2C**). *Rop,* is involved in vesicle release and control of synaptic activity (6)(7). In contrast, comparing all cancer lists with the downregulated embryonic RNAs in the *Top3β* mutants, revealed 50 genes shared between the six different lists (**Figs. 2D, E**). Their KEGG pathway analysis indicated their involvement in the MAPK (mitogen-activated protein kinase) signaling pathway that regulates various cellular processes, including cell growth, proliferation, differentiation, and survival (8) (9). Thus, it seems that the MAPK signaling pathway is particularly sensitive to *Top3β* inactivation.

We also compared the features of the Drosophila RNAs shared between the data sets in **Figs. 2A, B** and compared them with the ones present only in the human disease-derived list or the fly list of embryonic RNAs that depend on *Top3β* **(Supplementary Fig. S4**). With the exception of JME, the RNAs shared between the two datasets had on average longer 3’ and 5’ UTRs than the orthologs of the human disease RNAs that were not in the *Top3β*-dependent dataset. Similarly, in most cases, the pre-mRNAs in the overlapping group were significantly longer. This result is consistent with the human *Top3β* also affecting the expression of RNAs with the same features as its fly homolog, pointing to the potential value of identifying direct Top3β target mRNAs in the model system because their reduced expression might contribute to the disease phenotype.

### *In vivo* RNA interactions of Top3β in the absence of bulk transcription

By capturing mRNA targets directly during the first 2 hours of embryogenesis, we should be able to focus on direct RNA targets of Top3β and avoid indirect effects. Furthermore, because this time window ends before zygotic transcription takes off, we do not expect Top3β to bind to pre-mRNAs in this assay and we should be able to focus on cytoplasmic interactions. To identify directly the RNA targets to which Top3β binds and performs its topoisomerase activity, we sequenced the RNAs attached to Top3β::eGFP by immunoprecipitation (IP) in the presence of limited amounts of ionic detergents. This was done using animals that expressed an eGFP fused to wild-type Top3β and, as a negative control, Top3β*^Y332F^*::eGFP in which Y332 was replaced by a phenylalanine (Y332F). This mutant form of Top3β lacks only the -OH group through which Top3β forms the transient covalent bond between its Y332 side chain and the RNA target during the enzymatic reaction (10). We confirmed by qMS analysis that the wild-type and the mutant Top3β::eGFP fusions were expressed at similar levels **(Supplementary Table S9**). The naturally occurring covalent bond between Top3β and its targets makes artificial cross-linking superfluous (thus reducing the problem of high background) while allowing the use of ionic detergents at low concentrations during the isolation. These conditions should pull down primarily covalently bound RNAs without denaturing the antibodies and their interactions. Because we expected that only a small fraction of the potential targets would be bound at the time of tissue lysis, we tried to preserve the covalent bond between Top3β’s Y332 and the RNA targets for the IP by complexing Mg^2+^, which is needed for the enzyme to resolve the bond at the end of the enzymatic reaction (11)(90). To determine if the target RNAs were specifically bound to the –OH group of Y332, we used the Y332F mutant as a negative control as described above. The RNAs that are enriched in the Top3β^+^::eGFP (Top3β::eGFP) IP compared to the Y332F mutant are candidate targets of the catalytic activity of Top3β **(Supplementary Table S10)**. Because long RNAs seem more likely to depend on this enzymatic activity, we further tested the quality of the data by comparing the RNA length of the presumed targets with the background RNA length. Indeed, longer RNAs were more likely to covalently bind to Top3β as they showed higher enrichment in IPs from embryonic extracts containing Top3β::eGFP compared to the Top3β^Y332F^::eGFP control (**Fig. 3A**). It appears possible that a minor fraction of the suspected target RNAs represents false positives if their expression is upregulated by Top3β::eGFP but not by Top3β^Y332F^::eGFP expression. However, such an effect would not be expected to lead to the size correlation observed **(Fig. 3A)**. This argues that if such an effect exists, it would be only a minor one. The size correlation results might also reflect that longer RNAs have a higher chance of enduring physical stress and are, also for this reason, more likely to become substrates of the enzymatic activity of Top3β. Interestingly, recent work in human cells also found that TOP3β preferentially binds to long mRNAs (74).

**Fig 3:**
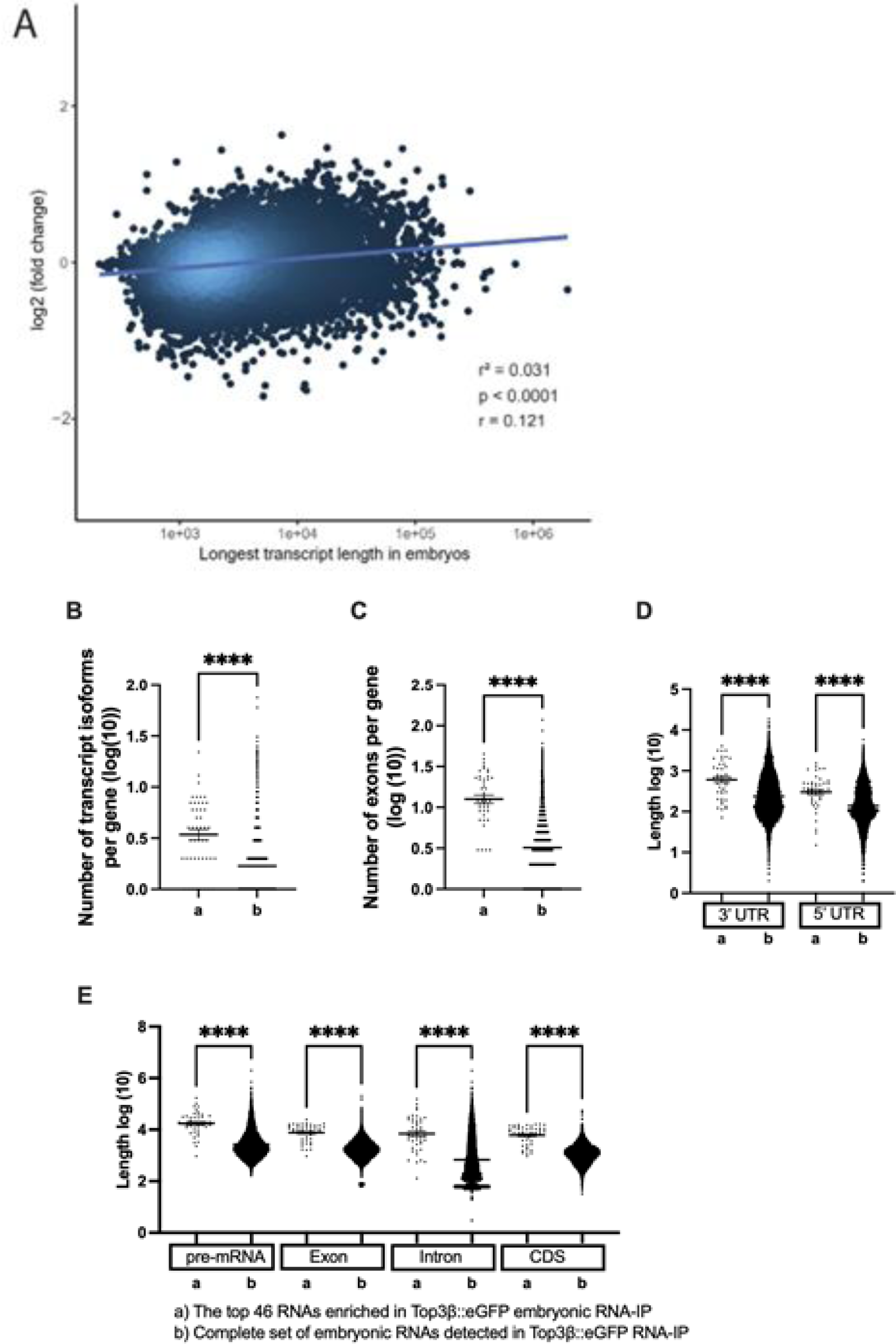
**Characteristics of embryonic RNA targets of Top3β::eGFP**. **A)** Longer RNAs are more likely targets of the enzymatic activity of Top3β because they are more enriched in the IP with the wild-type *Top3β* compared to the *Top3β^Y332F^*. r measures the strength of the correlation and p stands for p-value. **B-E)** Transcript features of the top 46 embryonic RNA targets of Top3β::eGFP. The complete sets of RNAs detected in the wild-type RNA-IP served as the standard. p-value <0.0001=****, p-value <0.001=***, p-value <0.01=**, p-value <0.05=*.

From the list of immunopurified RNAs, the ones with an adjusted p-value smaller than 0.05 (adjp-value<0.05) and a log2-fold change of more than 1 (log2FC>1) were selected and ranked based on their adjp-value. We used these parameters primarily because it is difficult to capture the covalently bound phase. We noted that *Rop* was close to the cut-off with an adjp-value of 0.027 and a log2-fold change of 0.54 (**Supplementary Table S10**). 46 embryonic RNAs and their genes fulfilled these criteria and were used for further analyses (**Supplementary Table S11**). The features that these RNAs share, can either indicate features that increase the chance of becoming a target of the enzymatic activity of Top3β or correlate with another feature that increases the chance of becoming a target. Drosophila embryonic Top3β RNA targets displayed on average a higher number of transcript isoforms and exons per gene, and longer 3’- and 5’UTRs compared to the complete set of different types of RNAs detected in the Top3β^+^::eGFP IPs (**Figs. 3B-D, Supplementary Tables S10-S11)**. The fact that targeted mRNAs have longer 3’UTRs is consistent with mRNAs that undergo localization and/or translational regulation having a higher chance of becoming a Top3β target because these processes are mainly controlled by sequences in the 3’UTR. A higher abundance of mRNAs with long coding sequences (CDS) might reflect that mRNAs with long open reading frames (ORFs) might become a Top3β target when their translation stalls (**Fig. 3E, Supplementary Tables S10-S11)**. Whereas the longer introns and the higher number of exons might indicate that mRNA splicing increases the chance of becoming a Top3β target, the absence of transcription in the first two hours of embryogenesis seems to argue that these mRNAs are more likely to become targets because of other criteria that also apply to this group of genes. In summary, these results present evidence for the involvement of Top3β with long and complex mRNAs and mRNAs that are localized and/or under translational control. This is also consistent with a reported finding that long mRNAs are overrepresented among human mRNAs artificially crosslinked to Top3β in tissue culture cell lines (74).

### Connecting the Drosophila Top3β data to mammalian Top3β, FMR1, and FMRP

The Drosophila proteins copurifying with Top3β::GFP support its suggested roles in translation, interacting with cytoplasmic RNAs and RNP granules (**Supplementary Table S9; Supplementary Fig. S5**). The interaction network shows that one branch of its interactions goes through Tdrd3 and FMR1, the fly homolog of the fragile X mental retardation syndrome protein (FMRP) (**Supplementary Fig. S5B**). Mammalian Top3β has been linked to FMRP, too, and it binds to ribosomes and regulates translation (1). These researchers also found an overlap between HITS-CLIP data produced with FMRP and mouse brain extracts and Top3β HITS-CLIP data obtained with Flag-Top3β and HeLa extracts. Comparing the Drosophila Top3β::eGFP RNA-IP results with the shared mammalian result set, revealed that homologs of 12 of the 46 top maternal embryonic Drosophila targets were also present in the combined mammalian list. Considering that data not only from different species but also from different tissues were compared, this strong overlap reveals mRNA targets that support mainly cellular physiology in different cell types (**FIG. 4A**). These results further strengthen the value of the technique applied to obtain the Drosophila Top3β target RNAs and the use of Drosophila as a model to study the role of Top3β in human disease. The common feature of the proteins encoded by the shared RNA targets is that they are predominantly large proteins. Many are localized to the plasma membrane or the cell cortex (e.g., Sev, Mgl, α-Spec, Kst, Pcx) or cytoskeleton-associated (e.g., Dhc64C, Mhc, Myo10A, α-Spec, Kst), have axonal functions (e.g., Shot, Dhc64C, Kst, Prosap), or scaffolding functions (Shot, Prosap). Consistent with similar results in vertebrates (74), these results show that there is a clear overlap between the Top3β and the FMRP activities but, additionally, the two proteins might perform activities independently of the other one.

We also compared the effects of Drosophila *Top3β* and *FMR1* on total RNA levels for each gene in 0-2 hrs old embryos. For this, we used the results from the *Top3β^26^* null mutant (Adjp <0.0002; log2-fold change <-1 and >1, respectively; **Supplementary Table S1**) and the published results from an *FMR1* mutant (92). These analyses consider reads per gene and do not discriminate between different RNA isoforms. Consistent with their physical interaction, more than half of the genes affected in the *FMR1* mutant were also found affected in *Top3β^26^* (**Fig. 4B**). In both cases, only a few genes showed higher RNA levels in the mutant and the overlap between the two elevated RNA datasets was very low. Lists of the genes identified in the datasets of both mutants are presented in the supplements (**Supplementary Table S13**).

**Fig. 4:**
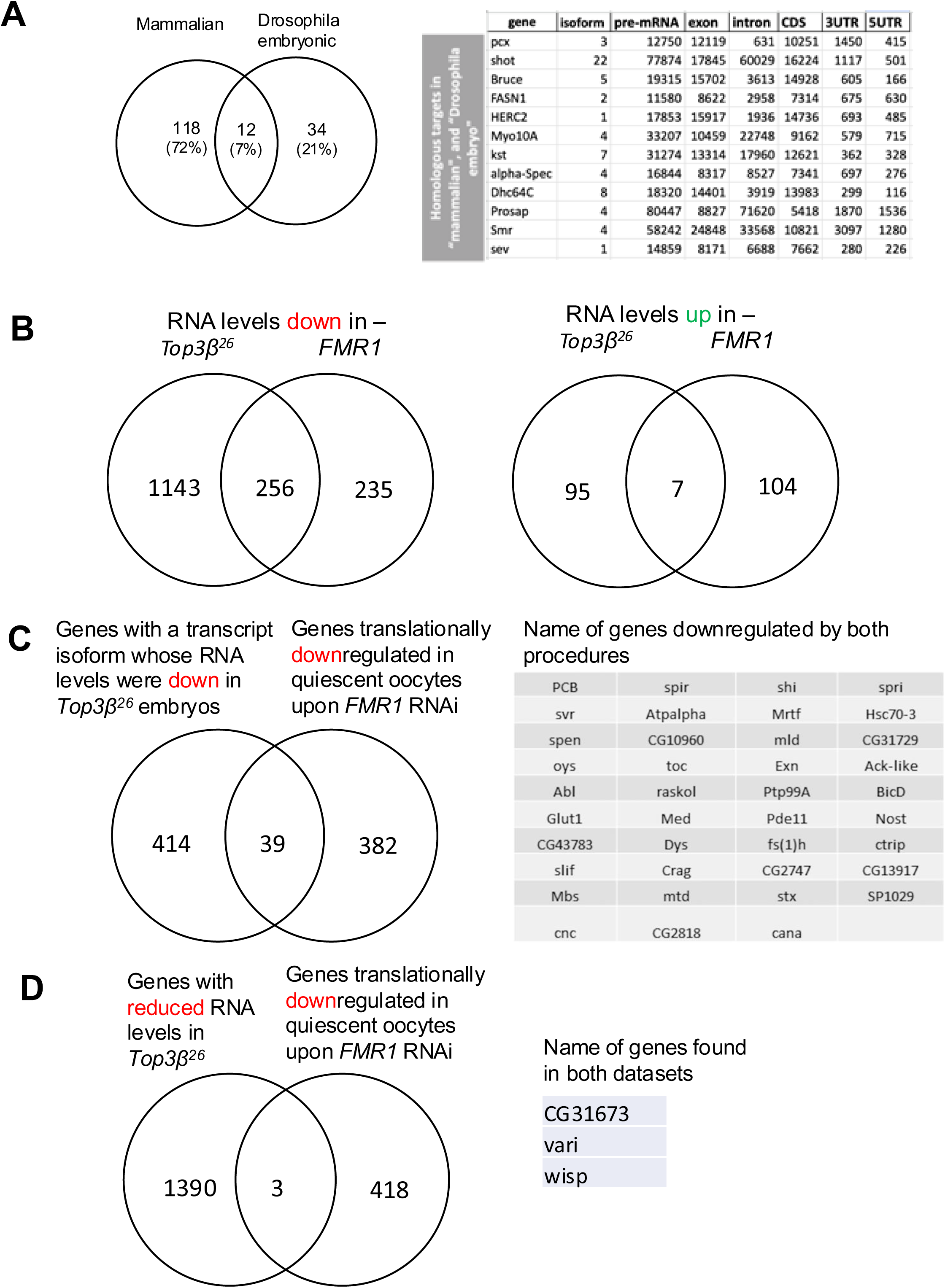
Comparing Drosophila Top3β targets and effects to the corresponding FMR1/FMRP and mammalian Top3β data. **A)** A previous mammalian study produced a list of overlapping mouse FMRP targets in brain extracts and Top3β targets in HeLa cells (1). The Venn diagram compares the list of Drosophila homologs of these mammalian targets to our list of direct targets of Top3β. The list of the Drosophila homologs that were also found in both mammalian interactor screens is shown with the characteristics of the Drosophila gene. The length measurements stem from the longest pre-mRNA isoforms (**Supplementary Table S11**). **B)** Comparing RNASeq data from 0-2 hrs old embryos from the *Top3β^26^* null mutant (Adjp < 0.0002; log2-fold change <-1 and >1, respectively, according to **Supplementary Table S1)** and the *FMR1* mutant (92). The analysis considers reads per gene. Lists of the genes identified in the overlapping datasets are presented in the supplements **(Supplementary Table S13). C)** Genes encoding transcript isoforms that depend on *Top3β* for their normal accumulation in young embryos (**Supplementary Table S4**) and transcripts that depend on *FMR1* for their translation in quiescent oocytes (48). The Venn diagram shows the overlap between the two datasets and the genes in the overlap are listed. **D)** Comparing RNASeq data from 0-2 hrs old *Top3β^26^*embryos (Adjp <0.0002; log2-fold change <-1**; Supplementary Table S1)** and the *FMR1* RNAi effect on translational repression (48).

The role of Drosophila *Fmr1* in translation was also studied in quiescent oocytes using RNAi treatment and analysis in ovaries (48). Comparing the list of their genes significantly repressed in translation to the transcript isoforms less abundant in the extracts of *Top3β^26^*mutants (**Supplementary Table S4**) revealed several shared mRNAs (**Fig. 4C)**. Even though different tissues had been used for the two experiments and different effects had been compared, there is still an overlap between the two data sets. Again, *FMR1* and *Top3β* seem to act on many of the same mRNAs, but they also seem involved in additional functions, independently of each other. We also compared the effect of lack of Top3β on the RNASeq reads in 0-2 hrs old embryos per gene with the effect the RNAi treatment against *FMR1* has on translational repression (48) in the ovary. Using the data from **Supplementary Table S1** and the same conditions as in **Fig. 4B** (Adjp <0.0002; log2-fold change <-1) we only found three hits in the overlap with the same *FMR1* data (**Fig. 4D**), which had shown a good overlap with the list of reduced transcript isoforms. Two reasons seem to explain the lower number in this overlap compared to the ones in **B)** and **C)**. One, as mentioned by the authors, *FMR1* RNAi knockdown did not generally reduce mRNA levels except for its own RNA (48). In contrast, the *FMR1* mutant used in the study used for **B**) (92), reduced many RNA levels. Two, studying the effect on transcript isoforms (as in **C)** is more sensitive to detecting level changes because the same changes might be judged insignificant in the pool of different transcript isoforms of the same gene.

### Top3β supports the expression and subcellular localization of its transcript targets

The *Top3β^Y332F^*and *Top3β^26^*mutants have reduced or no function, but the mutations cause neither lethality nor sterility. This is a common phenomenon observed with conserved genes and proteins, which are still important for optimizing cellular physiology. In a lab setting, the reduced function of these mutants can often be compensated for by other mechanisms. However, by directly analyzing the cellular phenotypes of the mutants, one may observe increased cellular stress and changes in physiology or anatomy that reveal the cellular function of the normal gene. Such cellular stress and alteration often lead to signs of premature aging, like a decline in neuromuscular performance and a shorter lifespan.

To evaluate the consequences of the interaction of Top3β with its RNA targets, we selected *shot* (encoding a spectraplakin), *Dhc64C* (encoding the large subunit of the cytoplasmic dynein motor), and *kst* (encoding the β_Heavy_-spectrin) from the top list of RNAs covalently bound to Top3β::eGFP for further analysis. Aside from exhibiting the typical RNA features of the top targets, their homologs were also identified by the mammalian studies used for **Fig. 4A** (1) and they are expressed in neurons, too (22)(23). In normal young embryos, *shot* mRNA accumulates more strongly in the polar granules and the pole cells at the posterior end (**Fig. 5)**. In embryos that were maternally and zygotically mutant for *Top3β*, *shot* mRNA levels were overall reduced. Additionally, the more intense, localized staining pattern in the posterior pole region was lost in the null- and the enzymatic mutant *Top3β^Y332F^*, but not in *Top3β^ΔRGG^*, suggesting that the normal expression and localization of *shot* mRNA depends on *Top3β* and particularly on its enzymatic activity. Analogous studies revealed that the anti-Shot antibody signal was similarly weaker in *Top3β^26^* and *Top3β^Y332F^* embryos compared to the wild type (**Fig. 5)**, suggesting that the reduced mRNA levels caused the reduced Shot protein expression. These findings are consistent with *shot* mRNA being a primary target of Top3β and they highlight the importance of the enzymatic activity of Top3β for the efficient expression and localization of the encoded protein. *Top3β*^ΔRGG^ mutant embryos still show a reduction of the Shot protein signal even though the RNA is still localized **(Fig. 4)**. This is consistent with the role of Top3β in the translation of *shot* RNA. That Top3β is required for efficient translation of mRNAs has already been demonstrated (74). The effects on the proteins encoded by two more *Top3β* targets were similar. The cytoplasmic Dynein heavy chain (Dhc or Dhc64C) protein accumulated more strongly at the posterior end of the wild-type embryo, in front of the pole cells. This was neither detectable in the null mutant nor in *Top3β^Y332F^* **(Fig. 5)**. The Kst protein accumulated at the apical cortex in normal blastoderm embryos, but in the *Top3β* null mutant and *Top3β^Y332F^*, its signal was reduced throughout the embryo, including the apical cortex. Protein signals for Dhc showed an intermediate strength in the *Top3β^ΔRGG^* mutant embryos **(Fig. 5)**, indicating that the RGG box also affects their expression, albeit less strongly than the catalytic activity. We conclude that *Top3β* facilitates gene expression and particularly localized accumulation of proteins at sites distant from the nucleus.

**Fig. 5.**
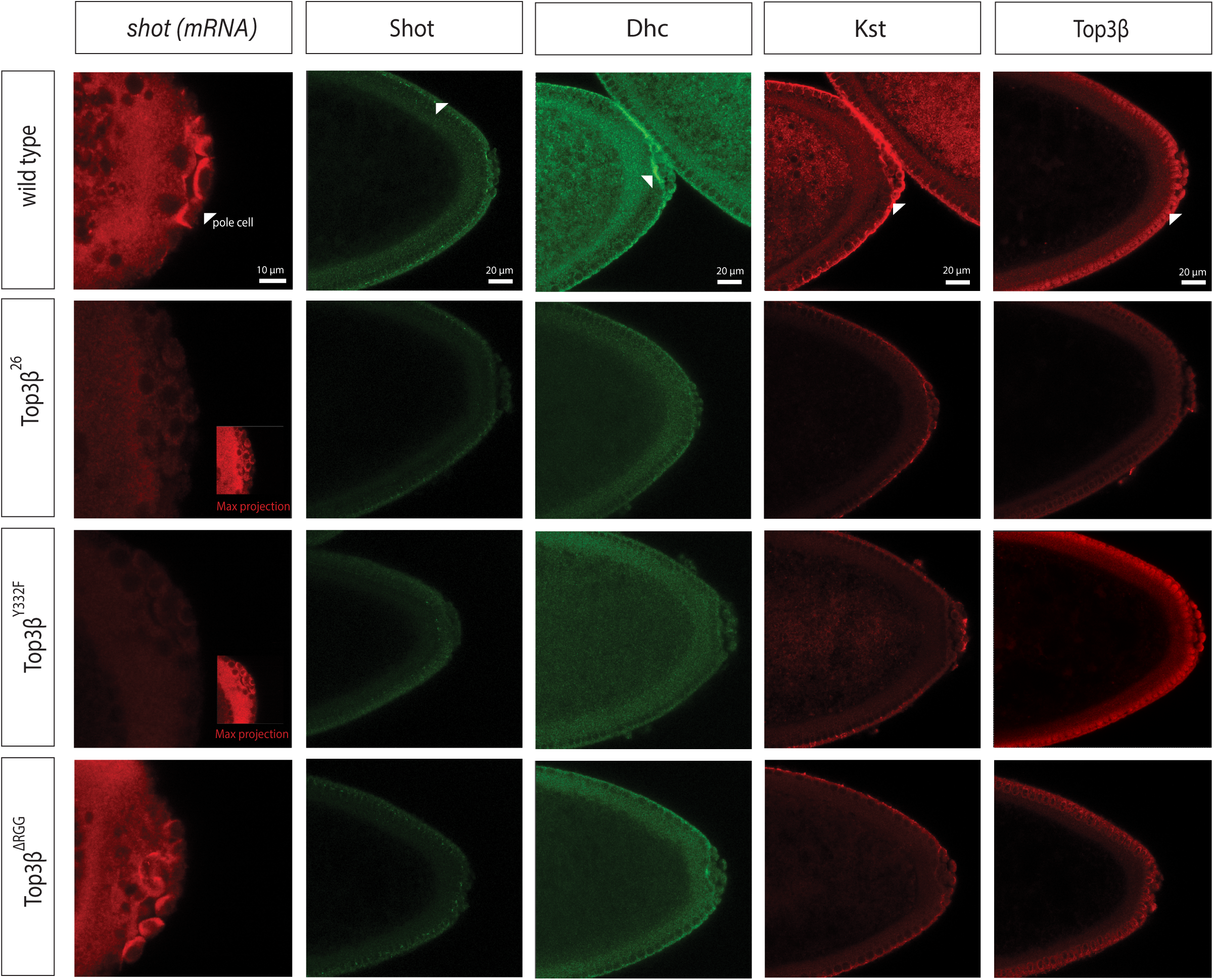
Expression and localization of gene products of Top3β targets. The posterior pole of the embryos is shown to the right. *shot* mRNA and Shot protein localization in the posterior part of syncytial (*shot* mRNA) and cellularizing (Shot protein) wild-type embryos in the different *Top3β* mutants. Lower signal levels and less localized signals were seen in the null and *Top3β^Y332F^*. Both were less reduced in *Top3β^ΔRGG^* mutant embryos. Note the *shot* mRNA localization in the pole cells in the wild type and *Top3β^ΔRGG^* mutant. Dhc64C localization at the posterior side of the somatic part of the embryo (arrowhead) was not detectable in the null mutant and *Top3β^Y332F^* and reduced in *Top3β^ΔRGG^*mutant embryos. Kst localization at the apical membrane (arrowhead) was reduced in null mutants and *Top3β^Y332F^* compared to the wild type. Top3β protein is localized similarly to its targets. It is, however, absent from the null mutant which reveals the background levels of the staining. Note that Top3β is also nuclear in the wild type and *Top3β^Y332F^* but absent from nuclei in the *Top3β^ΔRGG^* embryo. Imaging settings were identical for the staining for the same protein and mRNA, respectively, in the different genotypes except for the insets which were overexposed to reveal the embryo. The scale bar is 10µm for the *shot* mRNA pictures and 20µm for the other micrographs.

Top3β normally localizes to the same area in blastoderm embryos as the different proteins and mRNAs described here. In the Discussion section, we will detail how this seems to contribute to an interesting gene expression mechanism. Note also that the wild-type protein and Top3β^Y332F^ localize to the cytoplasm and the nuclei **(Fig. 5)**. In contrast, Top3β^ΔRGG^ is seen above background levels only in the cytoplasm. This indicates that its RGG box is needed to localize Top3β to the nucleus in blastoderm embryos and that the lack of the RGG box likely reduces the nuclear functions of Top3β_ΔRGG_.

### *Top3β* supports neuronal structure and function in aging adults

The negative geotaxis assay measures the activity of the nervous system and the muscles, and the coordination between them. In this assay, all four *Top3β* genotypes showed similar climbing abilities in young flies **(Fig. 6A).** Neuromuscular conditions sometimes become only apparent at an older age because the young nervous system can compensate for some cellular deficits. But such compensations can take their toll if they stress the cells. We, therefore, tested for premature aging and age-related degeneration of the climbing ability. Indeed, as seen in **Fig. 6A**, from week 6 on, the *Top3β^Y332F^* and *Top3β^26^* flies became slower, and their climbing ability declined more than the wild-type one. The results suggest that Top3β plays a role in aging and combating neurodegeneration in *Drosophila* motor neurons.

**Fig. 6:**
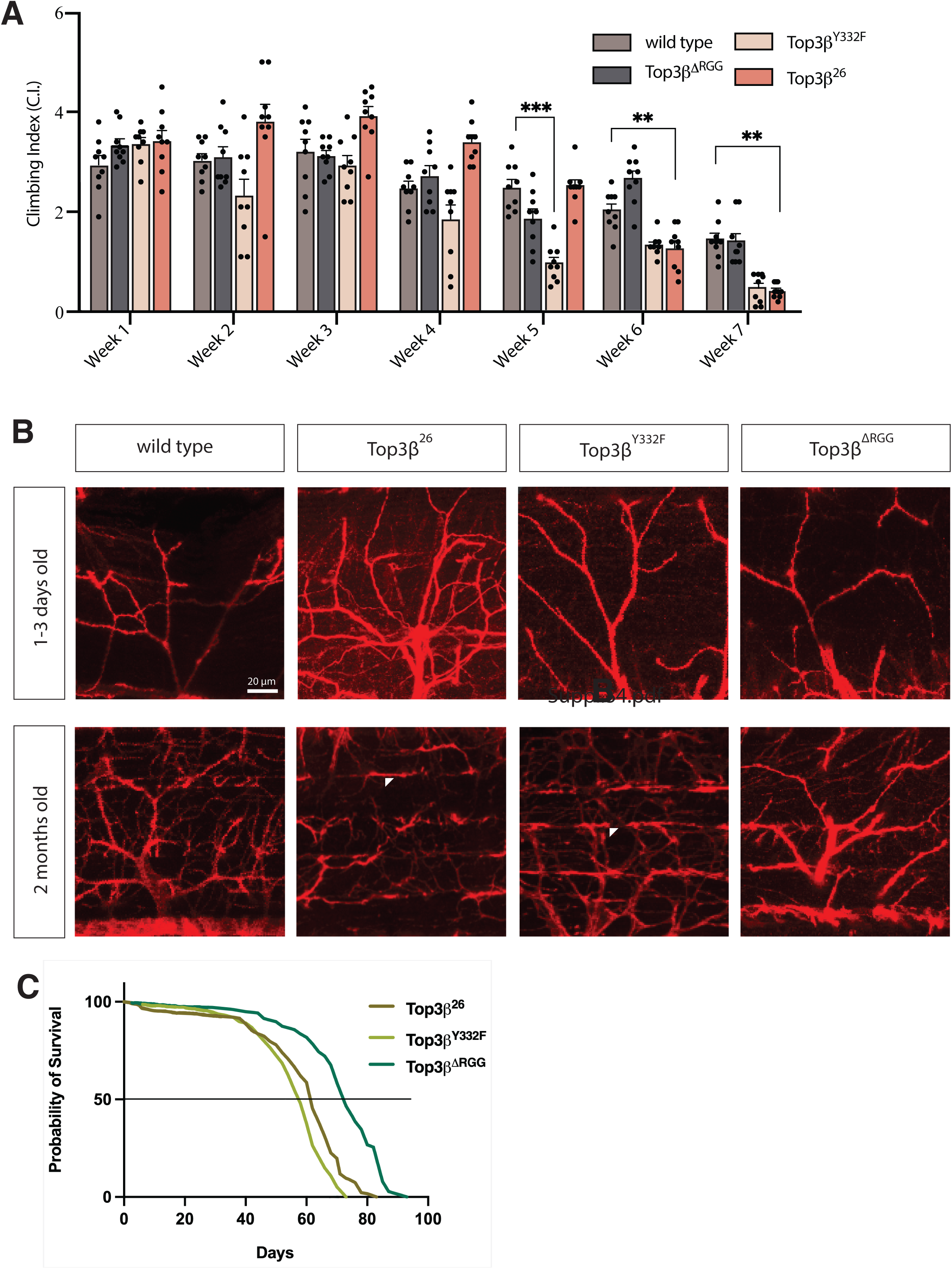
Aging effects of *Top3β* mutants. **A)** Climbing assays of wild type, *Top3β^ΔRGG^*, *Top3β^Y332F^*, and *Top3β^26^* flies. A two-way ANOVA analysis was performed. The graph bars show the mean climbing index (C.I.) for the different genotypes. Error bars represent the standard error of the mean (S.E.M). Each dot represents a calculated individual C.I. The adjusted p-values for each significant difference are indicated on the graph. (p-value <0.0001=****, p-value <0.001=***, p-value <0.01=**, p-value <0.05=*.) **B)** In adults, null mutant NMJs contain more branches than the wild type, indicative of a compensatory mechanism. Arrowheads point to the lost synaptic integrity of the dorsal longitudinal flight muscle (DLM). NMJ synapses were visualized by staining motor neurons with anti-HRP antibodies. The progressive denervation of the DLM is particularly evident in the null and the *Y332F* mutant in the adult thoracic NMJs. The scale bar represents 20µm. All panels show maximum intensity projections with a z-step size of 0.7μm. **C)** Life span shortening by *Top3β* mutations. *Top3β^26^*null and *Top3β^Y332F^* mutants reduce long-term survival. *Top3β^ΔRGG^*, which showed only a minor reduction compared to a wild-type strain in other experiments, served as a control because it was induced in the same background as *Top3β^Y332F^*.

In aging flies with declining motor abilities and neurotransmission, fragmentation of synaptic motor terminals becomes apparent (24). We studied the structure of the adult neuromuscular junctions (NMJs) in the four genotypes during aging **(Fig. 6B)**. Particularly the young null mutants contained more branched structures compared to the simpler branch structures in the wild type. This might constitute a mechanism to compensate for less efficient synapse function. Although to a lesser degree, young *Top3β^Y332F^* and *Top3β*^Δ*RGG*^ mutants also displayed presynaptic overgrowth. Concomitant with the premature loss of locomotive ability around week 6 **(Fig. 6A)**, denervation of the DLM (Dorsal Longitudinal Muscle) synapses became evident. Particularly in the null mutants, but also in *Top3β^Y332F^*, loss of synaptic integrity was observed during aging in 1-2 months old adult flies **(Fig. 6B)**. The micrographs of these mutants are likely an underestimate of the average effect of the mutation because *Top3β^26^*and *Top3β^Y332F^* flies live less long than *Top3β*^Δ*RGG*^ **(Fig. 6C)** and healthy flies, and only living flies were selected for the NMJ preparation. The decline of the NMJ structure and function was most likely more severe in the dead animals. In summary, it appears that the reduced expression of many Top3β targets can initially be compensated in neurons for some time, but over time they lead to premature aging of the nervous system and reduced neuromuscular performance. It should be pointed out that a shorter life span has also been reported for *Top3β^-/-^*mice (93).

### *Top3β* mutants enhance (G4C2)_49_-induced neuronal toxicity in flies

To gain deeper insights into the cellular significance of Top3β’s catalytic activity toward one specific RNA, we investigated its effect on the phenotype induced by GC-rich repeat RNA sequences. For this, we used the repeat sequence (GGGGCC)_49_ (also named G4C2)_49_ that induces neurotoxic effects in both humans and flies. In humans, the presence of 20-28 repeats of this sequence was shown to be associated with neurodegenerative diseases such as amyotrophic lateral sclerosis (ALS) and frontotemporal dementia (FTD) (25)(26)(27). Using a GMR-GAL4 driver (28) to study the effect of this RNA on the adult eyes revealed that *Top3β* mutants enhanced the neurotoxic effect of the (G4C2)_49_ transcripts in the eyes of young flies aged 1-2 days (**Fig. 7**). Of the *Top3β* mutants, *Top3β^26^* and *Top3β^Y332F^* exhibited the most severe eye lesions with irregular ommatidial structures, and this was particularly evident from the widespread cell death. Young *Top3β^ΔRGG^*mutants displayed irregular ommatidia but usually less cell death. After one month, the rough eye phenotype caused by the *Top3β^26^* and *Top3β^Y332F^* mutants affected the entire eye, and widespread degeneration of ommatidial cells was observed. These results show that the topoisomerase activity of Top3β counteracts the toxic activity of the (G4C2)_49_ transcripts in the Drosophila model for this disease **(Fig. 7)**.

**Fig. 7:**
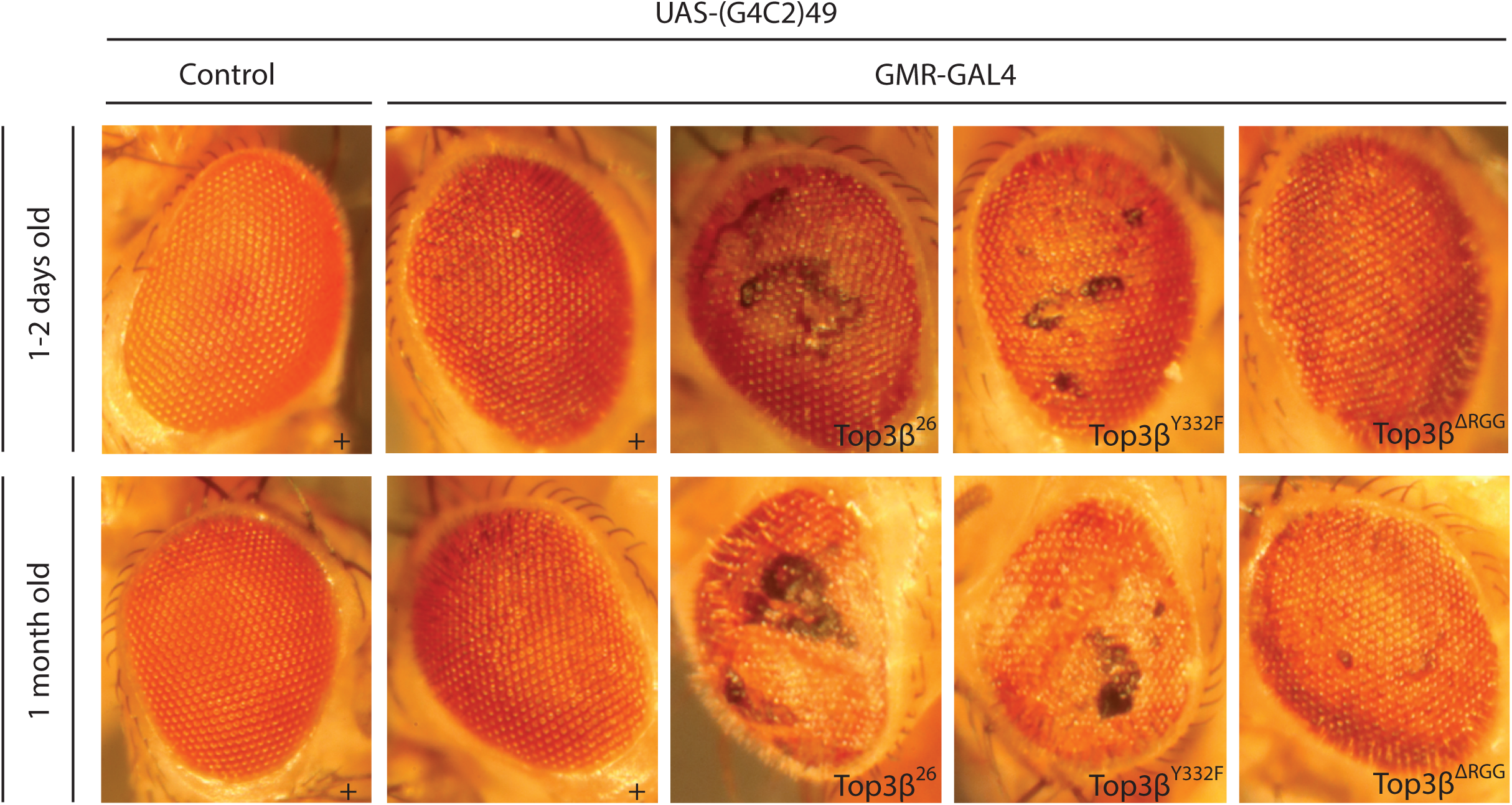
***Top3β* activities toward (G4C2)_49_ repeat RNAs in the eye.** Expression of the UAS- (G4C2)_49_ transgene in wild-type (+) and *Top3β* mutant flies using the GMR-GAL4 driver. *Top3β^ΔRGG^* caused mild disruptions, the *Top3β^26^* and *Top3β^Y332F^* mutants caused strong degeneration. Shown are data from one experiment carried out in parallel; data were reproduced in a total of three or more independent experiments. All *Top3β* mutants enhanced the degenerative phenotype of the (G4C2)_49_ RNA. UAS-(G4C2)_49_ and GMR-GAL4 lines were heterozygous. + indicates the control genotype *Top3β^+^*; the mutant alleles are indicated.

## Discussion

Top3β and its catalytic activity have a strong effect on the production of normal expression levels of a considerable number of RNAs during embryonic development. The features of these RNAs are the same as found in RNAs that are direct targets of the topoisomerase activity of Top3β. This points to important cytoplasmic functions of the Top3β topoisomerase in facilitating the translation of RNAs, translational control, and RNA localization. For instance, longer ORFs increase the risk of stalling translation because structural problems of mRNAs arise more frequently and prevent them from entering the ribosome. Top3β can rescue such mRNAs (74). The top Top3β mRNA targets depended not only on *Top3β* for their normal expression levels, but also for their localization within the cell (**Fig. 5**). Consistent with the proposed role of Top3β in rescuing RNAs with structural problems, the absence of the RNA topoisomerase activity does only reduce levels of affected RNAs and does not abolish them. Additionally, we also found evidence for nuclear roles in splicing, alternative splicing, and 3’end formation. The protein interaction map (**Supplementary Fig. S5**) supports these interpretations as well as its function in RNAi (49).

RNA localization combined with translational control is a mechanism to provide local expression of gene products. It is most important in large cells, but also for fine-tuning gene expression in general. From our results, it seems likely that this lack of fine-tuning gene expression of many genes, particularly the ones with the characteristics of *Top3β* targets and *Top3β*-dependent transcripts, makes human *Top3β* mutations a risk factor for several neurological diseases and cancers.

### RNA targets of Top3β

The Top3β interaction with splicing factors **(Supplementary Fig. S5; Table S9)** seems to explain why Top3β’s RNA targets often have a greater number of exons, longer introns, and alternative transcripts **(Fig. 3)**. These parameters might require additional RNA topoisomerase activity in the nucleus where Top3β presumably interacts with splice factors and the pre-mRNA. As with many other splice factors, Top3β might remain associated with the spliced mRNA in the cytoplasm (30)(31)(32)(33)(34). Nuclear translocation of Top3β requires the RGG motif in the blastoderm embryo **(Fig. 5)**. Therefore, the *Top3β*^Δ*RGG*^ allele should be a useful tool to analyze the nuclear localization and functions of Top3β. The RGG motives can be arginine methylated in HeLa cells (89), where it was shown that this enhances the interaction with TDRD3, the topoisomerase activity, and stress granule localization upon Arsenite treatment. Whether and how this methylation affects the nuclear localization of Top3β is, however, not clear yet.

The entanglement of long mRNAs and pre-mRNAs poses a considerable problem during transport over extended distances. Some of these processes involve additional mRNA packaging and unpackaging before the mRNAs can be activated for translation at the right place and time (94). The resulting structural problems might then interfere with their transport or, later, with their translation on ribosomes. The coding sequence (CDS) and untranslated regions (UTRs) of mRNA molecules are important for regulating gene expression at the level of RNA localization and translation. Particularly 3’UTRs affect not only the translation in neurons but mostly also the localization to the places where these mRNAs will be translated in response to extrinsic stimuli, such as neurotrophins, synaptic activation, and regeneration after injury (36)(8)(37)(38) (94). Consistent with the proposed function of Top3β in rescuing (pre-)mRNAs with such functions and characteristics, the *in vivo* targets of Top3β displayed a strong bias for these characteristics (**Fig. 3; Supplementary Table S11**) and the enzymatic activity of *Top3β* is particularly important for mRNAs that are generated from long pre-mRNAs, for mRNAs with long 3’UTRs and CDSs (**Figs. 1B-D; Supplementary Table S7**).

### Important targets of Top3β

*Rop*, encodes a Sec-1-like protein involved in vesicle trafficking in neuronal synapses and non-neuronal vesicle trafficking. In neuronal tissues, it regulates neurotransmitter secretion in a dosage-dependent manner (6)(7). In *Rop* mutants, the release of neurotransmitters from the pre-synapse area into the synaptic cleft fails to occur. In 7 neurological diseases with which human *Top3β* had been associated and in Drosophila embryos, normal expression levels of the *Rop* mRNA depend on *Top3β* (**Fig. 2C**; **Supplementary Table S7**)*. Rop* is likely a rewarding gene to investigate these neurological diseases. It was the only gene that was present in all 8 datasets and the *Rop* mRNA was a significant direct target of Top3β **(Supplementary Table S10).**

*shot* was identified in most experimental datasets analyzed. *shot* mRNA is a top Top3β target (**Supplementary Tables S10, S11**), in Drosophila and HeLa cells and also as an FMRP target in mouse brains (**Fig. 4A**; (1)(74)). In young Drosophila embryos, normal levels of a specific *shot* mRNA isoform also depend on *Top3β* and its enzymatic activity (**Supplementary Table S5**), and *shot* mRNAs and protein levels appeared reduced and less localized in embryos **(Fig. 5).** *shot* encodes a spectraplakin family member, a large cytoskeletal linker molecule that binds actin and microtubules and participates in axon growth and neuromuscular junction maintenance and growth control (39),(40),(41),(42). Drosophila *shot* has additional roles in many other tissues and cell types, and it consists of 45 exons producing 22 transcript isoforms, as opposed to 2 isoforms per gene, which is average for *Drosophila*. The reduced *shot* mRNA and protein levels and localization might reflect roles of Top3β in splicing *shot* pre-mRNAs, localizing *shot* mRNA, resolving RNA entanglements, and facilitating *shot* mRNA translation.

Localization of mRNAs combined with translational control is a mechanism that is particularly important in large cells (like oocytes and young embryos) and cells with large extensions (like neurons) (94). Many of the identified mRNAs function in such cells and have important roles in neurons because they encode large cytoskeletal and cortical elements (e.g., *kst*, *α-spec*, *shot, mgl*) or regulators of NMJ growth (e.g., *Prosap*). Along with α-Spec, Kst (β_H_-Spectrin) crosslinks F-actin and acts as a molecular scaffold when recruited to the apical membrane of epithelial cells (43)(44)(Fig. 5). Because cortical localization seems to be one of the processes that is most dependent on the RNA topoisomerase activity, it is relevant that Top3β accumulates preferentially at the apical cortex of embryos, too. Higher Top3β activity in this location is expected to contribute to the apical translatability of mRNAs – for instance by rescuing entangled mRNAs -, thereby expressing the encoded protein apically at higher levels. Therefore, the subcellular localization of Top3β activity seems to serve as an additional tool to target gene expression efficiently to specific cellular locations. Such a mechanism can even produce protein localization if the mRNA is expressed uniformly.

### Top3β environment and relevance

The enrichment of neuronal-, stress-, and other ribonucleoprotein (RNP) forming granules, was a clear result of the analysis of the Top3β interactors (**Supplementary Fig. S5; Supplementary Table S9**). In the case of the neuronal RNP granules, they concentrate specific sets of mRNAs and regulatory proteins in the same location, promoting their common long-distance transport to axons or dendrites (20)(21). They also have a dual function in regulating the translation of associated mRNAs (47). As a component of neuronal RNPs, Me31B mediates the transport and controls the translation of neuronal RNAs, including the translational repression of synaptic transcripts (26). We found Me31B highly enriched with Top3β and this interaction depended on the RGG box, suggesting that the RGG box can link Top3β to neuronal RNPs. It will be interesting to find out whether this localization is also controlled by Arg-methylation of these motives as reported for stress granule localization of Top3β (89).

The lack of RNA topoisomerase activity can interfere with the expression of many different genes involved in a variety of different pathways. The resulting suboptimal functioning of different cellular mechanisms is likely to interfere with cellular physiology, making it less efficient and producing cellular stress. Further research needs to test which of the individual pathways suggested by our work causes a physiological problem in the human conditions involving *Top3β*. Serious neurological health issues often arise when several pathways are not functioning properly anymore, making it difficult to pinpoint a single primary cause. Our work revealed that reduced activity of Top3β affects several of the pathways that rely on mRNAs that are Top3β targets. The combination of defects in different physiological pathways is likely to impact the outcome of the various neurodevelopmental and cognitive disorders and cancers with altered *Top3β* function.

## Material and Methods

### Fly stocks and maintenance

The *white* (*w)* strain was used as a wild-type control as this was the genetic background of the mutants used in this study. The mutant fly stocks *Top3β^Y332F^* and *Top3β^ΔRGG^* were established as described below. *Top3β^26^* was obtained from Lee and colleagues (49) and described by Wu et al (50). The fly stocks were grown and maintained on a regular corn medium (ingredients: corn, yeast powder, sucrose syrup, agar, potassium sodium tartrate tetrahydrate, water, nipagin, and propionic acid) supplied with brewer’s yeast for the experiments. All *Drosophila melanogaster* fly stocks were stored at 18°C in glass or plastic vials with day/night (12 h/12 h) light cycles. All experiments were performed at 25°C. (G4C2)_49_ flies (Stock #84727; (28)) express 49 pure 5’GGGGCC3’ repeats under the control of UAS. These repeats are neurotoxic and model the G4C2 repeats seen in human *C9orf72* that are associated with Amyotrophic Lateral Sclerosis and Frontotemporal Dementia.

### Generation of *Top3β* alleles

Genomic sequences were scanned for suitable sgRNA target sites with the JBrowse from FlyBase and appropriate sgRNA sequences were chosen based on (a) their distance from the tagging site, and (b) the absence of potential off-target sites on the same chromosome as the target site (51). To mutate the endogenous *Top3β*, a sgRNA was designed, which hybridized in the vicinity of the Y332 codon of *Top3β* and induced the Cas9 nuclease to perform a double-strand break. Complementary oligodeoxynucleotides with suitable overhangs (**Supplementary Table S12**) were annealed and cloned into the pCFD5 vector, which is used to express the sgRNA (52). The Cas9 fly stock was obtained from the Bloomington Stock

Center (#79004). ssDNA was designed for the mutation changing the Tyr (Y) to a Phe (F) codon and injected into the progeny produced by Cas9- and sgRNA-expressing flies. The single-strand oligodeoxynucleotide with the flanking arms precisely matching the endogenous sequences changes the sequence at the mutation site by serving as a template (**Supplementary Table S12**) for the homologous recombination DNA repair at the double-strand break. The same strategy was used to delete the endogenous RGG region in *Top3β* (βRGG) using the appropriate gRNA and ssDNA templates. Stocks were established from individual mutants and genomic sequencing confirmed the precise mutation events in the initial mutant fly and the final stock. Stocks were deposited in the Bloomington Drosophila Stock Center (presently as #99728 and #99728). Both point mutants produce normal RNA levels (**Supplementary Table S1-3**) and produce stable proteins **(Fig. 5)** – although Top3β^1′RGG^ was absent from the syncytial nuclei. That the point mutations of these two mutants do not affect protein levels was also shown by qMS analysis of the tagged versions of these proteins which gaave very similar results **(Supplementary Table S9**).

### RNA sequencing

RNA was isolated using 1 ml of TRIzol® Reagent per 50-100 mg of tissue samples (0-2 hours embryos). The samples were homogenized at room temperature. RNA extraction was done according to the Trizol protocol. The University of Bern’s Next Generation Sequencing Platform conducted quality control assessments, library generation, sequencing runs, and sequencing quality control steps. The purified total RNA was assessed for quantity and quality using a Thermo Fisher Scientific Qubit 4.0 fluorometer with the Qubit RNA BR or HS Assay Kit (Thermo Fisher Scientific, Q10211 or Q32855, respectively) and an Agilent Advanced Analytical Fragment Analyzer System using a Fragment Analyzer RNA Kit (Agilent, DNF-471), respectively. rRNA depletion and construction of sequencing libraries were performed according to Lexogen’s guidelines using a CORALL Total RNA-Seq Library Prep Kit with UDI 12nt Set A1 (Lexogen, 117.96). The libraries were paired-end sequenced using either shared Illumina NovaSeq 6000 SP, S1, or S2 Reagent Kits (200 cycles; Illumina, 200240719, 2002831, or 20028315) on an Illumina NovaSeq 6000 instrument. The quality of the sequencing runs was assessed using the Illumina Sequencing Analysis Viewer (Illumina version 2.4.7), and all base call files were demultiplexed and converted into FASTQ files using Illumina bcl2fastq conversion software v2.20.

The quality and quantity of reads generated from RNA-seq were evaluated. Most of the reads were from mature transcripts without introns, but we mapped them to the reference genome that includes introns using an alignment tool like Hisat2 that can handle large gaps. This required information on where each gene is located in the genome, available for example from Ensemble (53). The number of reads mapping to each gene was counted using the FeatureCounts tool. This resulted in a table of read counts for each sample and gene. The bulk RNA seq data has been posted on NCBI-GEO (https://www.ncbi.nlm.nih.gov/geo/query/acc.cgi?acc=GSE249429).

To test for differential expression between two experimental groups containing three biological replicates each, we used the DESeq2 tool. The analysis included the following steps: 1. Normalization to correct for differences in the total number of reads between samples. 2. Variance estimation between replicates to account for the limited number of replicates in RNA-seq experiments (DESeq2 incorporates information from other genes with similar overall expression levels into the estimation). 3. Log-fold change (LFC) adjustment based on evidence strength which was estimated by the LFC. If it is weak (e.g., because the gene is expressed at low levels, the variance between replicates is high, or there are few replicates), the LFC shrinks toward zero. 4. Calculation of a test statistic and comparison to the normal distribution to obtain a p-value. 5. Multiple test corrections using the Benjamini-Hochberg procedure to consider only potentially detectable, differentially expressed genes. DESeq2 applies a false discovery rate correction based on the Benjamini-Hochberg procedure. However, the multiple-test correction considers only genes that could potentially be detected as differentially expressed. Only these genes will have an adjusted p-value. The mean read count across all samples is used to decide if a gene should be included or not. Mapping of the sequence reads to specific isoforms was performed with the Salmon method (91). The Venn diagrams in Fig. 1A were produced with the online tool of the Ghent University (https://bioinformatics.psb.ugent.be/webtools/Venn/).

### Extracting transcript features

The information about exon length, exon count, 3’UTR length, gene name, gene length, 5’UTR length, mRNA length, and CDS length was extracted from the gtf file of dmel-all-filtered- r6.42.gff using a custom python script.

### Generation of the *Top3β::eGFP* cDNA clone for GAL4>UAS expression

For the *Top3β::eGFP* construct, the *Top3β* cDNA clone LD10035 was amplified. Enhanced GFP (EGFP) was PCR amplified from a plasmid (pEGFP–C3). PCR amplified fragments were run on an agarose gel and purified using the Wizard SV kit (Promega). The sequences were cloned into the pUASz1.0 vector (54) with the *eGFP* ORF at the C-terminal end of the *Top3β* ORF using a ligation kit (NEB). The UASz vector allows one to induce the constructs in any tissue, including the somatic tissue, the germline, and the CNS (54). Plasmids were sequenced to confirm the integrity of the construct and purified with a PureYield kit (Promega) for injection. Transgenic stocks were established with the attP landing platform 86F from the Bloomington Stock Center (#24749) and also deposited in the Stock Center (#99726).

To generate an enzymatically dead enzyme, a Phe codon substituted the codon for the Tyr responsible for crosslinking the enzyme with the nucleic acid in the *UASz-Top3β::eGFP* vector to give rise to *UASz-Top3β^Y332F^::eGFP*. Similarly, to figure out the role of the RGG box, we deleted in the *UASz-Top3β::eGFP* vector the sequence coding for this box, giving rise to *UASz- Top3β^1′RGG^::eGFP*. At the site of the desired mutation, oligonucleotide primers with the desired mutation were used to amplify the mutant double-stranded DNA plasmid (**Supplementary Table S12)**. A protocol based on two different PCR amplifications was used for this. First, two elongation reactions were carried out, with one of the two primers each. In the second reaction, the two initial elongation reactions were mixed and an additional PCR reaction was carried out. The eGFP fusion constructs were transformed into *Drosophila* using the attP-86F landing platform. They are also available from the Bloomington Stock Center (#99725 and #99727). mat-tub-GAL4 (#7062) was used to drive the expression in the older female germline. The wild-type and the mutant eGFP fusions were expressed at similar levels as determined by qMS analysis **(Supplementary Table S9**). UASz-eGFP transgenic flies, obtained from the Bloomington stock center, were used as negative controls.

### Sample preparation for mass spectrometry

Drosophila Top3β::eGFP was purified using the GFP tag and a mouse anti-eGFP antibody as described previously (55). Protein G magnetic beads were washed 3x with PBS and incubated for 2 hours with an anti-GFP antibody. After this, they were washed 3x with 1 ml 1x PBS and one more time with 1 ml homogenization buffer (25 mM Hepes PH 7.4; 150 mM NaCl; 0.5 mM EDTA PH 8.0, 1 mM DTT, protease inhibitors). To prepare the extracts, 1 ml homogenization buffer was added to 1 gr of the samples (dechorionated embryos 0-2 hours) in the glass homogenizer. The samples were homogenized and transferred into the tube.

Homogenized samples were centrifuged at 16,000 g (RCF), 4°C for 40 min. The interphase was transferred into a fresh Eppendorf tube, centrifuged again at 16,000 g, 4°C for 40 min, and transferred into a fresh tube. Beads were then incubated with the extracts for 6 hours at 4°C on a wheel and the extracts were subsequently removed from the beads. The beads were washed once with 1 ml homogenization buffer and then several times with wash buffer (25 mM Hepes PH 7.4; 150 mM NaCl; 0.5 mM EDTA PH 8.0; 1 mM DTT; protease inhibitors). The wash buffer was removed, and the beads were sent for mass spectrometry.

### Mass spectrometry

Mass spectrometry analysis was done by the Proteomics and Mass Spectrometry Core Facility of the University of Bern, and the data are available in **Supplementary Table S9**. Proteins in the affinity pull-down were re-suspended in 8M Urea / 50 mM Tris-HCl pH8, reduced for 30 min at 37°C with 0.1M DTT / 100 mM Tris-HCl pH8, alkylated for 30 min at 37°C in the dark with IAA 0.5M / 100 mM Tris-HCl pH8, dilute with 4 volumes of 20 mM Tris-HCl pH8 / 2 mM CaCl2 before overnight digestion at room temperature with 100 ng sequencing grade trypsin (Promega). Samples were centrifuged, and a magnet holder trapped the magnetic beads to extract the peptides in the supernatant. The digests were analyzed by liquid chromatography (LC)-MS/MS (PROXEON coupled to a QExactive mass spectrometer, ThermoFisher Scientific) with three injections of 5 μl digests. Peptides were trapped on a µPrecolumn C18 PepMap100 (5 μm, 100 Å, 300 μm×5 mm, ThermoFisher Scientific, Reinach, Switzerland) and separated by backflush on a C18 column (5 μm, 100 Å, 75 μm×15 cm, C18) by applying a 60 min gradient of 5% acetonitrile to 40% in water, 0.1% formic acid, at a flow rate of 350 ml/min. The Full Scan method was set with a resolution at 70,000 with an automatic gain control (AGC) target of 1E06 and a maximum ion injection time of 50 ms. The data-dependent method for precursor ion fragmentation was applied with the following settings: resolution 17,500, AGC of 1E05, maximum ion time of 110 milliseconds, mass window 2 m/z, collision energy 27, underfill ratio 1%, charge exclusion of unassigned and 1+ ions, and peptide match preferred, respectively.

The data were then processed with the software MaxQuant (56) version 1.6.14.0 against the UniProtKB(57) Drosophila 7228 database (release 2021_02) containing canonical and isoform entries, to which common contaminants were added. The following parameters were set: digestion by strict trypsin (maximum three missed cleavages), first search peptide tolerance of 15 ppm, MS/MS match tolerance of 20 ppm, PSM and protein FDR set to 0.01 and a minimum of 2 peptides requested per group. Carbamidomethylation on cysteine was selected as a fixed modification; the following variable modifications were allowed: methionine oxidation, deamidation of asparagines and glutamines, and protein N-terminal acetylation; a maximum of 3 modifications per peptide was allowed. Match between runs was turned on (match time windows 0.7 min) but only allowed within replicates of the same kind.

Peptides were normalized by variance stabilization (58), imputed and combined to form Top3 (59) intensities, and considered alongside MaxQuant’s Label-Free Quantification (LFQ) values.

Missing peptides, respectively LFQ intensities, were imputed by drawing values from a Gaussian distribution of width 0.3 centered at the sample distribution mean minus 2.5x the sample standard deviation, provided there were at most 1 non zero value in the group; otherwise, the Maximum Likelihood Estimation (60) was used. Differential expression was performed by applying the empirical Bayes test (61) between groups; significance testing was performed as described (62).

### RNA immunoprecipitation

Protein G magnetic beads were washed 2-3x with 1ml blocking buffer (20 mM Hepes, 150 mM KCl, 20% Glycerol, 0.5 % Tween, BSA (Biolabs), Heparin 0.2 mg/ml, SDS 0.1%, 1/2 pill EDTA free protease inhibitors). Beads were blocked with a blocking buffer for 2-3 hours at room temperature. Anti-GFP antibody was added to the beads and rotated for 1-2 hours. Beads were washed 3x with non-hypotonic lysis buffer (20 mM Hepes, 1 mM EDTA, 150 mM KCL, 1 mM DTT, 0.5% Tween, 20% glycerol, SDS 0.1%, 1/2 pill EDTA free protease inhibitors).

Dechorionated embryos were 0-2 hours old. 1g of the embryos were homogenized in 2 ml hypotonic lysis buffer (20 mM Hepes, 10 mM KCL, 1 mM EDTA, 1 mM DTT, 0.5% Tween, SDS 0.1%, 1/2 pill EDTA free protease inhibitors, and RNase inhibitor (Biolabs, 40,000U/ml)). 600 μl of extract, 323 μl of Adjusting buffer (57% Glycerol 100%, 0.4M KCl, 1 mM EDTA, 1 mM DTT, 20 mM Hepes, SDS 0.1%), and 2 μl of RNase-free DNase I (20 μ/ml, Roche) were added to the washed antibody beads and were precipitated ON rotating at 4°C. Samples were washed 8x with 1 ml high salt wash buffer (20 mM Hepes, 1 mM EDTA, 200 mM KCl, 1 mM DTT, 0.5% Tween, 20% Glycerol, SDS 0.1%, 1/2 pill EDTA free protease inhibitors, and RNase inhibitor (Biolabs, 40,000U/ml)). On the last wash, 100 μl of the samples were taken before centrifugation. 100 μl of proteinase K buffer, 1.5 μl proteinase K (20 mg/ml Roche), and RNase inhibitors were added to each tube and were incubated for 30 minutes at 55°C. 1 ml Trizol reagent was added to each tube, and RNA was extracted according to the Trizol protocol (TRIzol™ Reagent, Invitrogen™, Catalog number: 15596026).

The Next Generation Sequencing Platform, University of Bern, conducted quality control assessments, library generation, sequencing runs, and sequencing quality control steps. The quantity and quality of the purified total RNA were assessed using a Thermo Fisher Scientific Qubit 4.0 fluorometer with the Qubit RNA BR or HS Assay Kit (Thermo Fisher Scientific, Q10211 or Q32855, respectively) and an Advanced Analytical Fragment Analyzer System using a Fragment Analyzer RNA Kit (Agilent, DNF-471), respectively. Sequencing libraries were made using a CORALL Total RNA-Seq Library Prep Kit, Version 1 with UDI 12nt Set A1 (Lexogen, 117.96) according to Lexogen’s guidelines for this kit. For the library preparation, purified RNA was neither depleted for rRNA nor selected for poly(A+) RNA but used directly for reverse transcription with random primers. Pooled cDNA libraries were paired-end sequenced using either shared Illumina NovaSeq 6000 SP, S1, or S2 Reagent Kits (200 cycles; Illumina, 200240719, 2002831, or 20028315) on an Illumina NovaSeq 6000 instrument. The quality of the sequencing run was assessed using Illumina Sequencing Analysis Viewer (Illumina version 2.4.7) and all base call files were demultiplexed and converted into FASTQ files using Illumina bcl2fastq conversion software v2.20.

The quality of the RNA-seq reads was assessed using fastqc v0.11.9 (63) and RSeQC v4.0.0 (64). Unique molecular identifiers (UMI) were extracted with umi-tools v1.1.2 (65), and adapters were trimmed with cutadapt v3.4.1 (66) according to manufacturer instructions. The reads were aligned to the Drosophila reference genome (BDGP6.32) using hisat2 v2.2.1 (67). FeatureCounts v2.0.1 (68) was used to count the number of reads overlapping with each gene as specified in release 103 of the Ensembl genome annotation. The Bioconductor package DESeq2 v1.36.0 (69) was used to test for differential gene expression between the experimental groups. To generate the log2 fold change versus transcript length plots, the estimated log2 fold change per gene was plotted against the most extended transcript length per gene with R v4.2.1 (70). We used the log2 fold change>=1 and the adjusted p-value (adjp-value) <0.05 of enrichment over control for further analysis.

### Immunolocalization and FISH

For the immunostaining of adult Dorsal Longitudinal Muscles (DLMs), thoraces of 0-3 days old and 2 months old adult wild-type*, Top3β^Y332F^*, *Top3β^ΔRGG^*, and *Top3β^26^* flies were separated from the rest of the body in 1x PBS. The thoraces were fixed in PFA (32% formaldehyde diluted to 4% with 1x PBS) for 30 min at room temperature and then washed 3x with PBS. All 1x PBS was removed from the tubes using a Pasteur pipette. The tubes were submerged into a liquid nitrogen flask for 10 seconds using cryogenic tweezers. Then, 300 μL of ice-cold 1x PBS was added to the samples. The thoraxes were cut into two hemithoraxes, transferred into 1x PBS (24), and blocked in 1x PBS with 0.1% normal goat serum and 0.2% Triton X-100 at pH 7.4 for at least one h at 4°C. Mouse anti-HRP or anti-Brp (nc82; 1:125 dilution) from Developmental Studies Hybridoma Bank were used to stain motor neurons. Primary antibodies were removed on the following day and the tissue was washed four times with PBST for 5 min each on a rotator. For the secondary antibody staining, goat anti-mouse A647 (1.5 mg/ml, diluted 1:200 dilution; Jackson ImmunoResearch Laboratories Inc.) was added to the samples, which were then kept at room temperature for 2 hours in a dark box on the rotator. After the 2 h incubation, the secondary stain was washed out 4 times for 5 min with PBST. Then, the samples were mounted on a slide (24).

Embryonic RNA in situ hybridizations (FISH) and antibody staining techniques, respectively, were carried out as described previously in references (81) and (82), respectively. Anti-βH- Spectrin/Kst antibody (no. 243 used at 1:500–1:600) was obtained from (83), anti-Shot (mAbRod1) and anti-Dhc (2C11-2) antibodies were obtained from the Developmental Studies Hybridoma Bank (https://dshb.biology.uiowa.edu).

### Negative geotaxis (climbing) assays

Wild-type flies and the three *Top3β* mutant strains were reared on fresh food and allowed to hatch at 25° C. For each genotype, all 0-3-day-old adult male flies were collected. Ten young adult male flies, each, were anesthetized and placed in 4 separate graduated flat-bottom glass cylinders. After 1 hour of recovery from CO_2_ exposure, flies were gently tapped 4x to the bottom of the cylinder to startle them repeatedly with constant force to displace the flies to the bottom surface. The flies were then allowed to climb the wall until they reached the top of the column. The climbing procedure was video recorded using an iPhone 11 Pro. These assays were repeated every week over 7 weeks on the same weekday. Three fly batches, each, provided the material for the triplicates which were performed within 5-minute intervals. After the three trials, each batch of 10 flies was transferred into glass vials and kept at 25° C. Before the experiment, every single batch was inspected to ensure that all 40 flies could be video-tracked when they started to climb after the tapping. To calculate each climbing index (CI), flies that remained at the bottom were counted as in zone 0. The formula used to calculate the CI is CI = (0 x n0 + 1 x n1 + 2 x n2 + 3 x n3 + 4 x n4 + 5 x n5) / nTotal, where nTotal is the total number of flies and nx is the number of flies that reached zone x.

### Statistics

All statistical analyses were performed using GraphPad Prism 8. For the negative geotaxis assays, the means, standard error of the means, and p-values were obtained by two-way ANOVA with Tukey’s multiple comparison test.

## Supporting information

Descriptions for Supplementary Tables

## Acknowledgments

Our thanks go to P. Nicholson and the members of the University of Bern NGS platform for sequencing the many RNA samples and to the IBU group of R. Bruggmann, particularly, S. Oberhänsli, G. van Geest, and R. Dörig for their important support during the analyses of the sequencing data and for providing the proof of the correct mutations for the reviewers. We thank the University of Bern (Switzerland) Mass Spectrometry Center (PMSCF) members S. Braga Lagache and N. Buchs for performing the MS experiments and M. Heller and A-C. Uldry for the data analysis. We wish to acknowledge all our group members who gave their time and expertise to provide critical input and technical support. Special thanks go to P. Vazquez, R. Dörig, and D. Beuchle. We highly appreciate the continuous support from friends and colleagues in the Drosophila community for sharing their results, fly stocks, and antibodies. For this work, too, FlyBase provided invaluable support, the Developmental Studies Hybridoma Bank provided antibodies, and the Bloomington Drosophila Stock Center (NIH P40OD018537) fly stocks.

## Funding

This work was supported by the Swiss National Science Foundation project grants 31003A_173188 and 310030_205075 and by the Equipment grant 316030_150824 to BS. Support came also from the University of Bern to BS. The funders had no role in study design, data collection, and interpretation, or the decision to submit the work for publication.

---

**Fig. S1:**
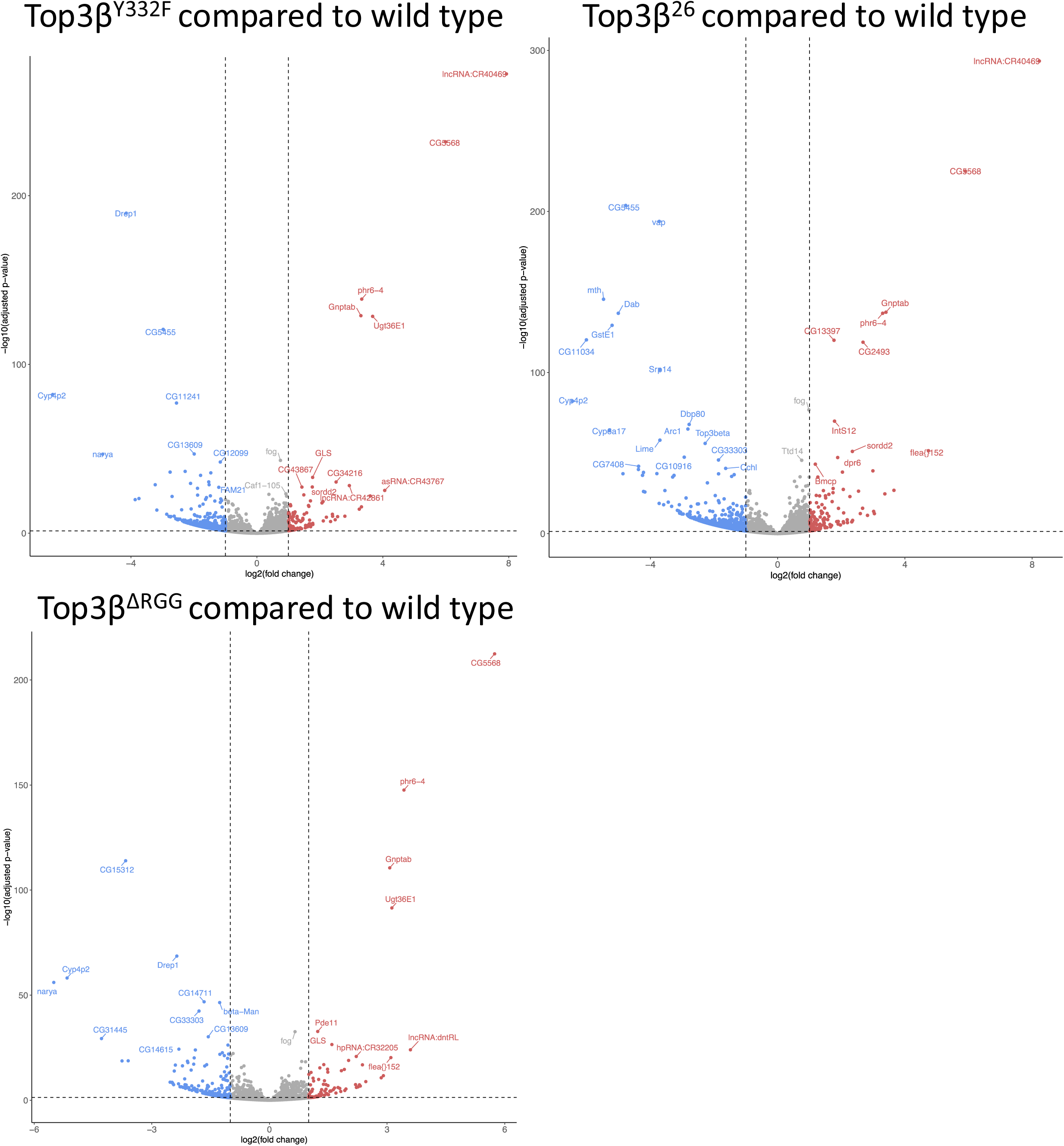
Transcriptomics of 0-2 hrs embryos. Volcano plots showing the effect of three *Top3β* mutations on the transcript levels in 0-2 hrs old *Drosophila* embryos. Adjusted p values are plotted against differences in transcript abundance. The logarithmic scales used for both axes are indicated.

**Fig. S2:**
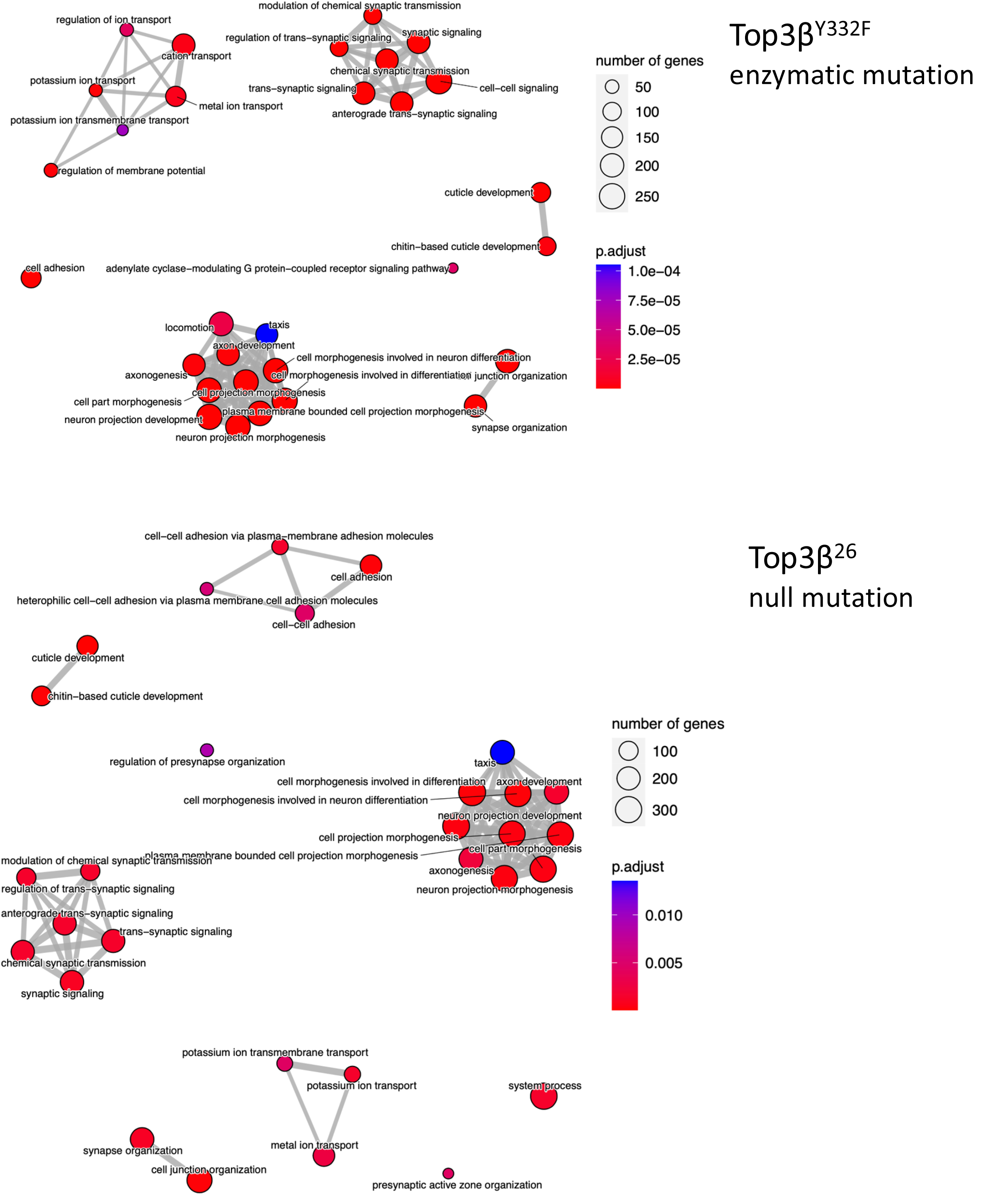
Gene Ontology Biological Processes. The enzymatic dead mutant *Top3β^Y332F^* and the null mutant *Top3β^26^*show very similar effects on biological processes, indicating that the Tyr in the active site plays a crucial role in the expression of normal levels of specific mRNAs at the early embryonic stage.

**Fig. S3:**
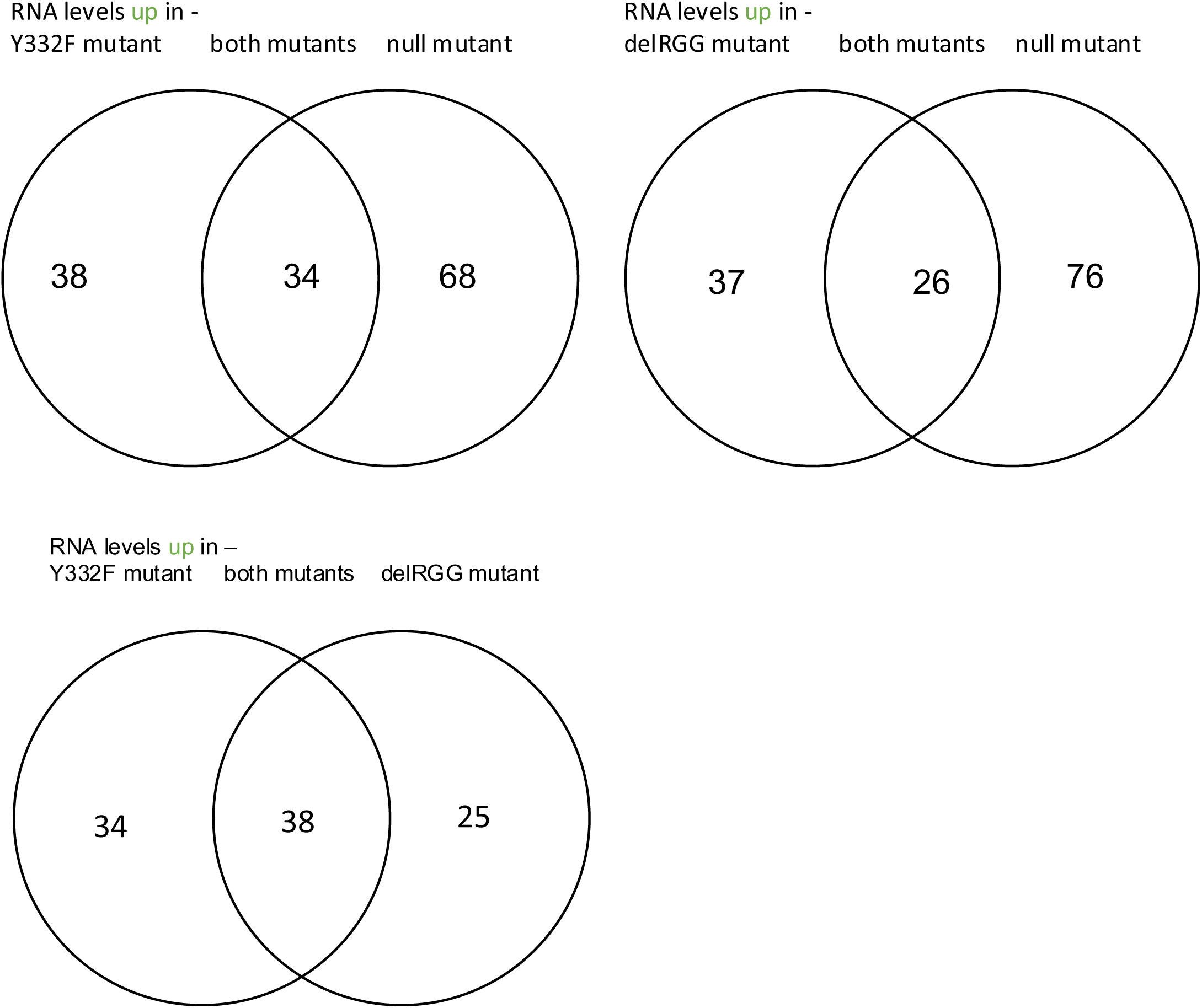
RNA levels elevated in the *Top3β* mutants. Genes showing higher transcript levels in 0-2 hrs old embryos mutant for *Top3β* (compared to their wild-type expression). Adjp <0.0002; log2 fold changes ≥1. The Venn diagram shows the pairwise overlap between the different mutants.

**Supplementary Fig. S4:**
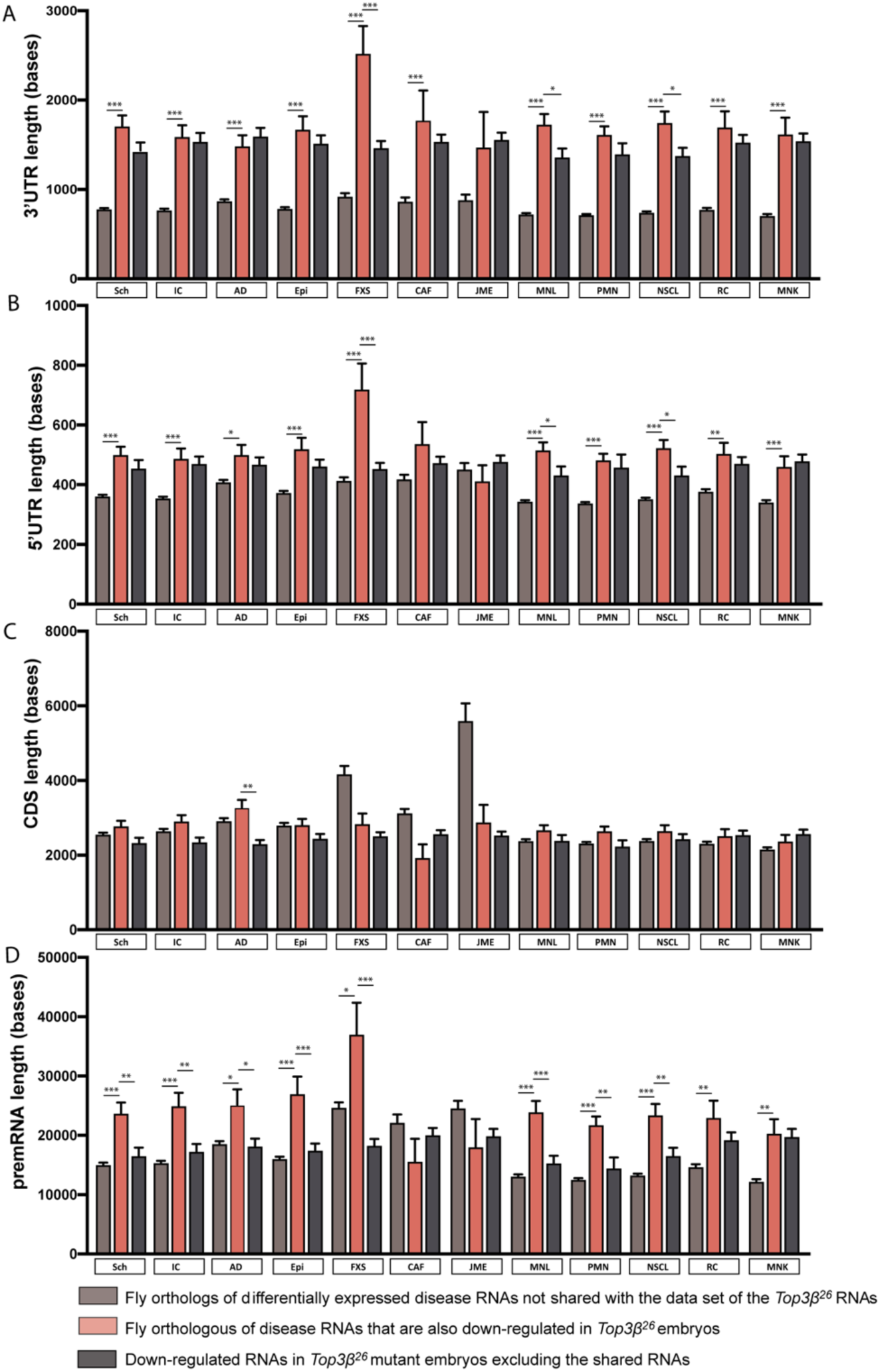
Transcript features for the differentially expressed RNAs from the embryo and the fly homologs of genes associated with the indicated diseases. A, B) mRNAs shared between the two lists show longer UTRs compared to the average fly homolog of the disease mRNA and the average *Top3β* dependent mRNA. C) CDS length revealed no significant difference between the three gene lists for each disease (except for AD). D) Shared mRNAs show longer pre-mRNAs compared to the average disease-homolog mRNA length and the average length of the embryonic *Top3β*-dependent RNA list. p-value <0.0001=****, p-value <0.001=***, p-value <0.01=**, p-value <0.05=*. Sch: Schizophrenia, IC: Impaired cognition, AD: Autistic Disorders, Epi: Epilepsy, FXS: Fragile X syndrome, CAF: Congenital anomaly of face, JME: Juvenile Myoclonic Epilepsy, MNL: Malignant neoplasm of the lung, PMN: Primary malignant neoplasm, NSCL: Non-Small Cell lung carcinoma, RC: Renal carcinoma, MNK: Malignant neoplasm of the kidney.

**Fig. S5:**
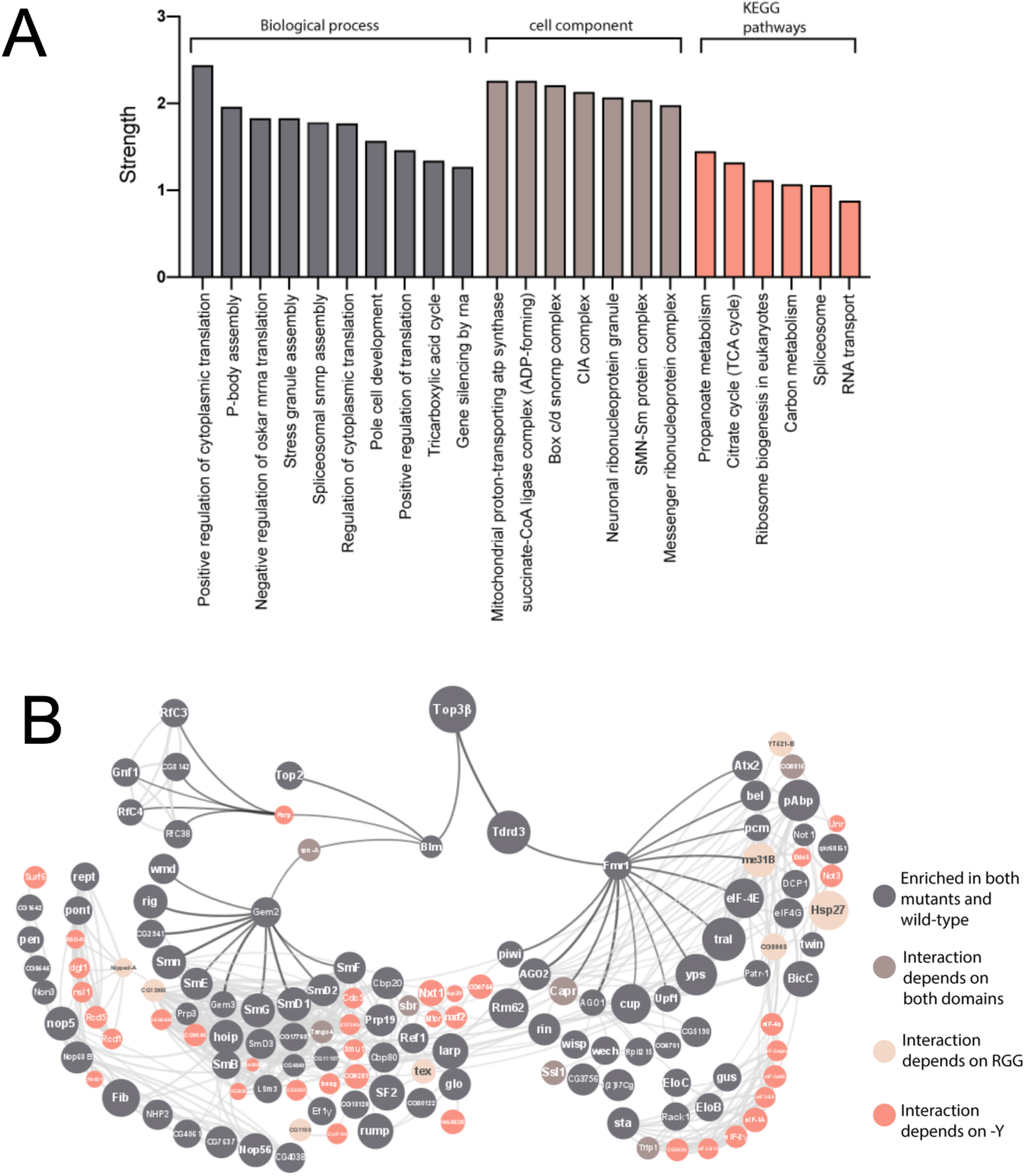
Pathways and interactions of Drosophila Top3β. A) *Gene ontology enrichment for Top3β-associated proteins for the top 100 interactors identified in embryonic extracts.* Immunoprecipitations (IP) on extracts from 0-2 hours old embryos were performed using *Top3β-eGFP* and the *eGFP* control line. Polypeptide components of the IP complexes were analyzed using mass spectrometry (MS) and their abundance was compared to the eGFP control. This resulted in a list of 426 potential complex components with an adjusted p-value <0.01 and log2FC>1 (**Supplementary Table S5).** 89 ribosomal proteins were in this set. A gene ontology enrichment analysis was then performed for the top 100 (according to log2FC) non-ribosomal proteins. This revealed proteins involved in the activation of translation, P-bodies, stress granules, and neural ribonucleoprotein granules, all membrane-less structures involved in storing specific mRNAs during periods of stress or concentrating mRNAs and regulatory proteins (12). Additionally, the interactors were also enriched in several functions related to mitochondria. B) *Proteins enriched in the embryonic protein-IP with their physical interactions according to Cytoscape-String.* Proteins that ended up further away from Top3β than the ones shown here, were removed from the interaction map. Tdrd3, an established interactor of Top3β (13), was among the top enriched proteins, suggesting that our immunoprecipitation was specific. To assess the role of the Y332 residue and the RGG box in Top3β interactions, IP results were compared with the ones from the *Top3β* mutants *Top3β^Y332F^-eGFP* and *Top3β^ΔRGG^-eGFP*. Among the 337 non-ribosomal binding candidates, 102 needed the Y residue and the RGG box to bind to Top3β*-*eGFP **(Supplementary Table S5)**. An additional 16 proteins needed the Tyr (but not the RGG) and 23 the RGG box (but not the Y332) for their binding to Top3β. A large fraction of the identified proteins is involved in RNA transport, translation, splicing, mRNA surveillance, and degradation. The physical interaction map of the proteins enriched in the Top3β::GFP IP was created using Cytoscape- String and the results from the mutant analyses were entered into this map. The resulting map resolved into two distinct sub-maps. The left branch connects Top3β functions to its nuclear functions and includes splicing factors. The right branch is enriched in cytoplasmic proteins and includes Tdrd3, FMR1, the piRNA pathway, and translation. The association first Tdrd3 and FMRP has already been described from work with vertebrate cell cultures (74). The interaction of Me31B with Top3β turned out to be dependent on the RGG box. As an RNA helicase, Me31B is a core component of a variety of ribonucleoprotein complexes (RNPs) that participate in translational control and mRNA decapping during embryogenesis, oogenesis, neurogenesis, and neurotransmission (14)(15)(16)(17)(18)(19)(20)(21). Me31B RNPs also containing Tral, eIF4E1, Cup, and pAbp. They are also involved in RNA localization and translational control of maternal mRNAs during oogenesis and embryogenesis (18). These other complex proteins were also enriched in the embryonic Top3β::eGFP IP.

## Notes

### Competing Interest Statement

The authors have declared no competing interest.

### Summary of Updates

Made improvements to text and Figures, clarified issues, added more data and supplements, and moved parts from Figures into supplements and the other way. Corrected mistakes.

## References

1. Xu D, Shen W, Guo R, Xue Y, Peng W, Sima J, et al. Top3β is an RNA topoisomerase that works with fragile X syndrome protein to promote synapse formation. Nat Neurosci. 2013;16(9):1238.

2. Ahmad M, Shen W, Li W, Xue Y, Zou S, Xu D, et al. Topoisomerase 3β is the major topoisomerase for mRNAs and linked to neurodevelopment and mental dysfunction. Nucleic Acids Res. 2017;45(5):2704–13.

3. Kaufman CS, Genovese A, Butler MG. Deletion of TOP3B is associated with cognitive impairment and facial dysmorphism. Cytogenet Genome Res. 2016;150(2):106–11.

4. Dekker NH, Rybenkov V V, Duguet M, Crisona NJ, Cozzarelli NR, Bensimon D, et al. The mechanism of type IA topoisomerases. Proc Natl Acad Sci. 2002;99(19):12126– 31.

5. Piñero J, Saüch J, Sanz F, Furlong LI. The DisGeNET cytoscape app: Exploring and visualizing disease genomics data. Comput Struct Biotechnol J. 2021;19:2960–7.

6. Wu MN, Littleton JT, Bhat MA, Prokop A, Bellen HJ. ROP, the Drosophila Sec1 homolog, interacts with syntaxin and regulates neurotransmitter release in a dosage-dependent manner. EMBO J. 1998;17(1):127–39.

7. Schulze KL, Littleton JT, Salzberg A, Halachmi N, Stern M, Lev Z, et al. Rop, a Drosophila homolog of yeast Sec1 and vertebrate n-Sect/Munc-18 proteins, is a negative regulator of neurotransmitter release in vivo. Neuron. 1994;13(5):1099–108.

8. Zhang W, Liu HT. MAPK signal pathways in the regulation of cell proliferation in mammalian cells. Cell Res. 2002;12(1):9–18.

9. Huang R, Quan Y, Chen J, Wang T, Xu M, Ye M, et al. Osteopontin promotes cell migration and invasion, and inhibits apoptosis and autophagy in colorectal cancer by activating the p38 MAPK signaling pathway. Cell Physiol Biochem. 2017;41(5):1851– 64.

10. Baker NM, Rajan R, Mondragón A. Structural studies of type I topoisomerases. Nucleic Acids Res. 2009;37(3):693–701.

11. Pommier Y, Sun Y, Shar-yin NH, Nitiss JL. Roles of eukaryotic topoisomerases in transcription, replication and genomic stability. Nat Rev Mol cell Biol. 2016;17(11):703.

12. Nostramo R, Xing S, Zhang B, Herman PK. Insights into the role of P-bodies and stress granules in protein quality control. Genetics. 2019;213(1):251–65.

13. Siaw GE-L, Liu I-F, Lin P-Y, Been MD, Hsieh T. DNA and RNA topoisomerase activities of Top3β are promoted by mediator protein Tudor domain-containing protein 3. Proc Natl Acad Sci. 2016;113(38):E5544–51.

14. Nakamura A, Amikura R, Hanyu K, Kobayashi S. Me31B silences translation of oocyte-localizing RNAs through the formation of cytoplasmic RNP complex during Drosophila oogenesis. 2001;

15. Wilhelm JE, Buszczak M, Sayles S. Efficient protein trafficking requires trailer hitch, a component of a ribonucleoprotein complex localized to the ER in Drosophila. Dev Cell. 2005;9(5):675–85.

16. Barbee SA, Estes PS, Cziko A-M, Hillebrand J, Luedeman RA, Coller JM, et al. Staufen- and FMRP-containing neuronal RNPs are structurally and functionally related to somatic P bodies. Neuron. 2006;52(6):997–1009.

17. Liu J, Carmell MA, Rivas F V, Marsden CG, Thomson JM, Song J-J, et al. Argonaute2 is the catalytic engine of mammalian RNAi. Science (80-). 2004;305(5689):1437–41.

18. Wang M, Ly M, Lugowski A, Laver JD, Lipshitz HD, Smibert CA, et al. ME31B globally represses maternal mRNAs by two distinct mechanisms during the Drosophila maternal-to-zygotic transition. Elife. 2017;6:e27891.

19. Lee J, Yoo E, Lee H, Park K, Hur J-H, Lim C. LSM12 and ME31B/DDX6 define distinct modes of posttranscriptional regulation by ATAXIN-2 protein complex in Drosophila circadian pacemaker neurons. Mol Cell. 2017;66(1):129–40.

20. Hillebrand J, Barbee SA, Ramaswami M. P-body components, microRNA regulation, and synaptic plasticity. ScientificWorldJournal. 2007;7:178–90.

21. Ruscica V, Bawankar P, Peter D, Helms S, Igreja C, Izaurralde E. Direct role for the Drosophila GIGYF protein in 4EHP-mediated mRNA repression. Nucleic Acids Res. 2019;47(13):7035–48.

22. Lee S, Kolodziej PA. Short Stop provides an essential link between F-actin and microtubules during axon extension. 2002;

23. Liu Z, Steward R, Luo L. Drosophila Lis1 is required for neuroblast proliferation, dendritic elaboration and axonal transport. Nat Cell Biol. 2000;2(11):776–83.

24. Sidisky JM, Babcock DT. Visualizing synaptic degeneration in adult Drosophila in association with neurodegeneration. JoVE (Journal Vis Exp. 2020;(159):e61363.

25. DeJesus-Hernandez M, Mackenzie IR, Boeve BF, Boxer AL, Baker M, Rutherford NJ, et al. Expanded GGGGCC hexanucleotide repeat in noncoding region of C9ORF72 causes chromosome 9p-linked FTD and ALS. Neuron. 2011;72(2):245–56.

26. Renton AE, Majounie E, Waite A, Simón-Sánchez J, Rollinson S, Gibbs JR, et al. A hexanucleotide repeat expansion in C9ORF72 is the cause of chromosome 9p21- linked ALS-FTD. Neuron. 2011;72(2):257–68.

27. Vatsavayai SC, Nana AL, Yokoyama JS, Seeley WW. C9orf72-FTD/ALS pathogenesis: evidence from human neuropathological studies. Acta Neuropathol. 2019;137:1–26.

28. Goodman LD, Prudencio M, Kramer NJ, Martinez-Ramirez LF, Srinivasan AR, Lan M, et al. Toxic expanded GGGGCC repeat transcription is mediated by the PAF1 complex in C9orf72-associated FTD. Nat Neurosci. 2019;22(6):863–74.

29. Mignone F, Gissi C, Liuni S, Pesole G. Untranslated regions of mRNAs. Genome Biol. 2002;3:1–10.

30. Ast G. How did alternative splicing evolve? Nat Rev Genet. 2004;5(10):773–82.

31. Black DL. Protein diversity from alternative splicing: a challenge for bioinformatics and post-genome biology. Cell. 2000;103(3):367–70.

32. Budagyan B, Loraine A. Gene length and alternative transcription in fruit fly. In: Proceedings 2004 IEEE Computational Systems Bioinformatics Conference, 2004 CSB 2004. IEEE; 2004. p. 515–6.

33. McGuire AM, Pearson MD, Neafsey DE, Galagan JE. Cross-kingdom patterns of alternative splicing and splice recognition. Genome Biol. 2008;9(3):1–19.

34. Fox-Walsh KL, Dou Y, Lam BJ, Hung S, Baldi PF, Hertel KJ. The architecture of pre-mRNAs affects mechanisms of splice-site pairing. Proc Natl Acad Sci. 2005;102(45):16176–81.

35. Liang J, Wen J, Huang Z, Chen X, Zhang B, Chu L. Small nucleolar RNAs: insight into their function in cancer. Front Oncol. 2019;9:587.

36. Han K, Gennarino VA, Lee Y, Pang K, Hashimoto-Torii K, Choufani S, et al. Human- specific regulation of MeCP2 levels in fetal brains by microRNA miR-483-5p. Genes Dev. 2013;27(5):485–90.

37. Tushev G, Glock C, Heumüller M, Biever A, Jovanovic M, Schuman EM. Alternative 3ʹ UTRs modify the localization, regulatory potential, stability, and plasticity of mRNAs in neuronal compartments. Neuron. 2018;98(3):495–511.

38. Glock C, Biever A, Tushev G, Nassim-Assir B, Kao A, Bartnik I, et al. The translatome of neuronal cell bodies, dendrites, and axons. Proc Natl Acad Sci. 2021;118(43).

39. Dewey EB, Parra AS, Johnston CA. Loss of the spectraplakin gene Short stop induces a DNA damage response in Drosophila epithelia. Sci Rep. 2020;10(1):1–13.

40. Sanchez-Soriano N, Travis M, Dajas-Bailador F, Gonçalves-Pimentel C, Whitmarsh AJ, Prokop A. Mouse ACF7 and drosophila short stop modulate filopodia formation and microtubule organisation during neuronal growth. J Cell Sci. 2009;122(14):2534–42.

41. Prokop A, Beaven R, Qu Y, Sánchez-Soriano N. Using fly genetics to dissect the cytoskeletal machinery of neurons during axonal growth and maintenance. J Cell Sci. 2013;126(11):2331–41.

42. Valakh V, Walker LJ, Skeath JB, DiAntonio A. Loss of the spectraplakin short stop activates the DLK injury response pathway in Drosophila. J Neurosci. 2013;33(45):17863–73.

43. Bennett V, Baines AJ. Spectrin and ankyrin-based pathways: metazoan inventions for integrating cells into tissues. Physiol Rev. 2001;81(3):1353–92.

44. Thomas GH, Williams JA. Dynamic rearrangement of the spectrin membrane skeleton during the generation of epithelial polarity in Drosophila. J Cell Sci. 1999;112(17):2843–52.

45. Banks GT, Fisher E. Cytoplasmic dynein could be key to understanding neurodegeneration. Genome Biol. 2008;9(3):1–4.

46. Bianco A, Dienstbier M, Salter HK, Gatto G, Bullock SL. Bicaudal-D regulates fragile X mental retardation protein levels, motility, and function during neuronal morphogenesis. Curr Biol. 2010;20(16):1487–92.

47. Formicola N, Vijayakumar J, Besse F. Neuronal ribonucleoprotein granules: Dynamic sensors of localized signals. Traffic. 2019;20(9):639–49.

48. Greenblatt EJ, Spradling AC. Fragile X mental retardation 1 gene enhances the translation of large autism-related proteins. Science (80-). 2018;361(6403):709–12.

49. Lee SK, Xue Y, Shen W, Zhang Y, Joo Y, Ahmad M, et al. Topoisomerase 3β interacts with RNAi machinery to promote heterochromatin formation and transcriptional silencing in Drosophila. Nat Commun. 2018;9(1):4946. DOI: 10.1038/s41467-018-07101-4

50. Wu J, Hou JH, Hsieh TS. A new Drosophila gene wh (wuho) with WD40 repeats is essential for spermatogenesis and has maximal expression in hub cells. Dev. Biol. 2006;296:219–230. doi: 10.1016/j.ydbio.2006.04.459.

51. Gratz SJ, Rubinstein CD, Harrison MM, Wildonger J, O’Connor-Giles KM. CRISPR-Cas9 genome editing in Drosophila. Curr Protoc Mol Biol. 2015;111(1):31–2.

52. Port F, Bullock SL. Augmenting CRISPR applications in Drosophila with tRNA-flanked sgRNAs. Nat Methods. 2016;13(10):852–4.

53. Cunningham F, Allen JE, Allen J, Alvarez-Jarreta J, Amode MR, Armean IM, et al. Ensembl 2022. Nucleic Acids Res. 2022;50(D1):D988–95.

54. DeLuca SZ, Spradling AC. Efficient expression of genes in the Drosophila germline using a UAS promoter free of interference by Hsp70 piRNAs. Genetics. 2018;209(2):381–7.

55. Vazquez-Pianzola P, Adam J, Haldemann D, Hain D, Urlaub H, Suter B. Clathrin heavy chain plays multiple roles in polarizing the Drosophila oocyte downstream of Bic-D. Development. 2014;141(9):1915–26.

56. Cox J, Mann M. MaxQuant enables high peptide identification rates, individualized ppb-range mass accuracies and proteome-wide protein quantification. Nat Biotechnol. 2008;26(12):1367–72.

57. Consortium U. UniProt: a worldwide hub of protein knowledge. Nucleic Acids Res. 2019;47(D1):D506–15.

58. Huber W, Von Heydebreck A, Sültmann H, Poustka A, Vingron M. Variance stabilization applied to microarray data calibration and to the quantification of differential expression. Bioinformatics. 2002;18(suppl_1):S96–104.

59. Silva JC, Gorenstein M V, Li GZ, Vissers JP. C; Geromanos, SJ Absolute quantification of proteins by LCMSE: A virtue of parallel MS acquisition. Mol Cell Proteomics. 2006;5:144–56.

60. Silver JD, Ritchie ME, Smyth GK. Microarray background correction: maximum likelihood estimation for the normal–exponential convolution. Biostatistics. 2009;10(2):352–63.

61. Kammers K, Cole RN, Tiengwe C, Ruczinski I. Detecting significant changes in protein abundance. EuPA open proteomics. 2015;7:11–9.

62. Uldry A-C, Maciel-Dominguez A, Jornod M, Buchs N, Braga-Lagache S, Brodard J, et al. Effect of Sample Transportation on the Proteome of Human Circulating Blood Extracellular Vesicles. Int J Mol Sci. 2022;23(9):4515.

63. Andrews S. FastQC: a quality control tool for high throughput sequence data. Babraham Bioinformatics, Babraham Institute, Cambridge, United Kingdom; 2010.

64. Wang L, Wang S, Li W. RSeQC: quality control of RNA-seq experiments. Bioinformatics. 2012;28(16):2184–5.

65. Smith T, Heger A, Sudbery I. UMI-tools: modeling sequencing errors in Unique Molecular Identifiers to improve quantification accuracy. Genome Res. 2017;27(3):491–9.

66. Martin M. Cutadapt removes adapter sequences from high-throughput sequencing reads. EMBnet J. 2011;17(1):10–2.

67. Kim D, Langmead B, Salzberg SL. HISAT: a fast spliced aligner with low memory requirements. Nat Methods. 2015;12(4):357–60.

68. Liao Y, Smyth GK, Shi W. featureCounts: an efficient general purpose program for assigning sequence reads to genomic features. Bioinformatics. 2014;30(7):923–30.

69. Love MI, Huber W, Anders S. Moderated estimation of fold change and dispersion for RNA-seq data with DESeq2. Genome Biol. 2014;15(12):1–21.

70. Core R. TEAM, 2017. R: A language and environment for statistical computing. R Foundation for Statistical Computing, Vienna, Austria. Online https://www.r-project.org. 2022;

71. Castells-Nobau A, Nijhof B, Eidhof I, Wolf L, Scheffer-de Gooyert JM, Monedero I, et al. Two algorithms for high-throughput and multi-parametric quantification of Drosophila neuromuscular junction morphology. J Vis Exp Jove. 2017;(123).

72. Szklarczyk D, Franceschini A, Wyder S, Forslund K, Heller D, Huerta-Cepas J, et al. STRING v10: protein–protein interaction networks, integrated over the tree of life. Nucleic Acids Res. 2015;43(D1):D447–52.

73. Stoll G, Pietiläinen OPH, Linder B, Suvisaari J, Brosi C, Hennah W, Leppä V, Torniainen M, Ripatti S, Ala-Mello S, Plöttner O, Rehnström K, Tuulio-Henriksson A, Varilo T, Tallila J, Kristiansson K, Isohanni M, Kaprio J, Eriksson JG, Raitakari OT, Lehtimäki T, Jarvelin MR, Salomaa V, Hurles M, Stefansson H, Peltonen L, Sullivan PF, Paunio T, Lönnqvist J, Daly MJ, Fischer U, Freimer NB, Palotie A. Deletion of TOP3β, a component of FMRP-containing mRNPs, contributes to neurodevelopmental disorders. Nat Neurosci. 2013 Sep;16(9):1228–1237. doi: 10.1038/nn.3484. Epub 2013 Aug 4. PMID: 23912948; PMCID: PMC3986889.

74. Su, S, Xue, Y., Sharov, A., Zhang, Y., Lee, S.K., Martindale, J.L., Li, W., Ku, W.L., Zhao, K., De, S., Shen, W., Sen, P., Gorospe, M., Xu, D., Wang, W. A dual-activity topoisomerase complex regulates mRNA translation and turnover. Nucleic Acids Res 2022; 50(12):7013–7033.

75. Schiavo G, Greensmith L, Hafezparast M, Fisher EM. Cytoplasmic dynein heavy chain: the servant of many masters. Trends Neurosci. 2013 Nov;36(11):641–51. doi: 10.1016/j.tins.2013.08.001.

76. Wilkie GS, Davis I. Drosophila wingless and pair-rule transcripts localize apically by dynein-mediated transport of RNA particles. Cell. 2001 Apr 20;105(2):209–19. doi: 10.1016/s0092-8674(01)00312-9. PMID: 11336671.

77. Bullock SL, Ish-Horowicz D. Conserved signals and machinery for RNA transport in Drosophila oogenesis and embryogenesis. Nature. 2001 Dec 6; 414(6864):611-6. doi: 10.1038/414611a. PMID: 11740552.

78. Foe, VE, and Alberts, BM. Studies of nuclear and cytoplasmic behavior in the five mitotic cycles that precede gastrulation in *Drosophila* embryogenesis. J. Cell Sci. 1983; 61, 31–70.

79. Lee, MT, Bonneau, AR, and Giraldez, AJ. Zygotic genome activation during the maternal-to-zygotic transition. Annu. Rev. Cell Dev. Biol. 2014; 30, 581–613

80. Tadros, W and Lipshitz, HD. The maternal-to-zygotic transition: a play in two acts. Development 2009; 136, 3033–3042.

81. Vazquez-Pianzola, P, Schaller, B, Colombo, M, Beuchle, D, Neuenschwander, S, Marcil, A, Bruggmann, R, and Suter, B. The mRNA transportome of the BicD / Egl transport machinery. RNA Biology 2017; 14:1, 73–89, DOI: 10.1080/15476286.2016.1251542

82. Vazquez-Pianzola, P, Beuchle, D, Saro, G, Hernández, G, Maldonado, G, Brunßen, D, Meister, P, and Suter, B. Female meiosis II and pronuclear fusion require the microtubule transport factor Bicaudal D. Development. 2022 Jul 1; 149(13):dev199944. doi: 10.1242/dev.199944.

83. Thomas, CM, and Kiehart, DP. β_Heavy_-spectrin has a restricted tissue and subcellular distribution during Drosophila embryogenesis. Development 1994; 120:7, 2039–2050. Doi: 10.1242/dev.120.7.2039

84. Rahman FU, Kim YR, Kim EK, Kim HR, Cho SM, Lee CS, Kim SJ, Araki K, Yamamura KI, Lee MN, Park SG, Yoon WK, Lee K, Won YS, Kim HC, Lee Y, Lee HY, Nam KH. Topoisomerase IIIβ Deficiency Induces Neuro-Behavioral Changes and Brain Connectivity Alterations in Mice. Int J Mol Sci. 2021 Nov 26;22(23):12806. doi: 10.3390/ijms222312806.

85. Zhu X, Joo Y, Bossi S, McDevitt RA, Xie A, Wang Y, Xue Y, Su S, Lee SK, Sah N, Zhang S, Ye R, Pinto A, Zhang Y, Araki K, Araki M, Morales M, Mattson MP, van Praag H, Wang W. Tdrd3-null mice show post-transcriptional and behavioral impairments associated with neurogenesis and synaptic plasticity. Prog Neurobiol. 2024 Feb;233:102568. doi: 10.1016/j.pneurobio.2024.102568.

86. Joo Y, Xue Y, Wang Y, McDevitt RA, Sah N, Bossi S, Su S, Lee SK, Peng W, Xie A, Zhang Y, Ding Y, Ku WL, Ghosh S, Fishbein K, Shen W, Spencer R, Becker K, Zhao K, Mattson MP, van Praag H, Sharov A, Wang W. Topoisomerase 3β knockout mice show transcriptional and behavioural impairments associated with neurogenesis and synaptic plasticity. Nat Commun. 2020 Jun 19;11(1):3143. doi: 10.1038/s41467-020-16884-4.

87. Su S, Xue Y, Lee SK, Zhang Y, Fan J, De S, Sharov A, and Wang W. A dual-activity topoisomerase complex promotes both transcriptional activation and repression in response to starvation. Nucleic Acids Research 2023, Vol. 51, No. 5 2415–2433. doi10.1093/nar/gkad086

88. Saha S, Sun Y, Huang SN, Baechler SA, Pongor LS, Agama K, Jo U, Zhang H, Tse-Dinh YC, Pommier Y. DNA and RNA Cleavage Complexes and Repair Pathway for TOP3B RNA- and DNA-Protein Crosslinks. Cell Rep. 2020 Dec 29;33(13):108569. doi: 10.1016/j.celrep.2020.108569.

89. Huang L, Wang Z, Narayanan N, Yang Y. Arginine methylation of the C-terminus RGG motif promotes TOP3B topoisomerase activity and stress granule localization. Nucleic Acids Res. 2018 Apr 6;46(6):3061–3074. doi: 10.1093/nar/gky103.

90. Yang X, Saha S, Yang W, Neuman KC, Pommier Y. Structural and biochemical basis for DNA and RNA catalysis by human Topoisomerase 3β. Nat Commun. 2022 Aug 9; 13(1):4656. doi: 10.1038/s41467-022-32221-3.

91. Patro, R., Duggal, G., Love, M. et al. Salmon provides fast and bias-aware quantification of transcript expression. Nat Methods 14, 417–419 (2017). 10.1038/nmeth.4197

92. Zhang, G., Xu, Y., Wang, X. et al. Dynamic FMR1 granule phase switch instructed by m6A modification contributes to maternal RNA decay. Nat Commun 13, 859 (2022). 10.1038/s41467-022-28547-7

93. Kwan, KY, and Wang, JC. Mice lacking DNA topoisomerase IIIbeta develop to maturity but show a reduced mean lifespan. Proc Natl Acad Sci USA 98(10):5717–21 (2001). DOI: 10.1073/pnas.101132498

94. Suter, B. RNA localization and transport. BBA - Gene Regulatory Mechanisms 1861; 10, 938–951 (2018). 10.1016/j.bbagrm.2018.08.004

