## Supplementary material for "RNA Targets and Physiological Role of Topoisomerase 3β": Descriptions for Supplementary Tables

**Supplementary Tables S1-S9**

**Supplementary Table S1)** lists the results of the transcriptome analysis of 0-2 hrs old embryos from *Top3β*$^{26}$ and compares it to the wild type.

**Supplementary Table S2)** lists the results of the transcriptome analysis of 0-2 hrs old embryos from *Top3β*$^{Y332F}$ and compares it to the wild type.

**Supplementary Table S3)** lists the results of the transcriptome analysis of 0-2 hrs old embryos from *Top3β*$^{\Delta RGG}$ and compares it to the wild type.

**Supplementary Table S4)** lists the results of the transcript isoforms that change their expression levels in *Top3β*$^{26}$ mutant embryos compared to the wild type.

**Supplementary Table S5)** lists the results of the transcript isoforms that change their expression levels in *Top3β*$^{Y332F}$ mutant embryos compared to the wild type.

**Supplementary Table S6)** lists the results of the transcript isoforms that change their expression levels in *Top3β*$^{\Delta RGG}$ mutant embryos compared to the wild type.

**Supplementary Table S7) 0-2 hrs embryonic transcript isoforms affected by mutations in** *Top3β.* This file lists the filtered Top3β transcriptomics results for specific mRNA isoforms (with adjusted p-value <0.05 and absolute log2 fold change >1). Columns C, D: Top3β$^{26}$ vs wild-type embryos. Columns E, F: Top3β$^{\Delta RGG}$ vs wild-type embryos. Columns G, H: Top3β$^{Y332F}$ vs wild-type embryos. Columns I: gene name (symbol), Columns J-M: length of indicated RNA features. The embryo background list (2$^{nd}$ sheet) shows the length of the RNA features for all transcripts identified.

**Supplementary Table S8). List of Drosophila genes whose human homologs are affected by the neurological diseases and cancers studied in Fig. 2.**

**Supplementary Table S9) List of proteins co-immunoprecipitated (co-IP) with Top3β::GFP.** Quantitative MS determined the enrichment in comparison to GFP-only IPs. co-IPs with mutant versions of Top3β*::GFP are also shown. They show the dependence of the interaction on the RGG domain and the Y332 residue.

**Supplementary Table S10) List of RNAs (and gene names) that were purified with Top3β::GFP from embryonic extract.** Enrichment of an RNA in the Top3β::GFP IP versus the Top3β$^{Y332F}$::GFP mutant protein IP was used to select the best target RNAs. RNA features are indicated for these RNAs, too.

**Supplementary Table S11) List of the top targets of Top3β::GFP** with their RNA features indicated. Length measurements shown are derived from the longest pre-mRNA isoforms.

**Supplementary Table S12) Oligos, primers, and fly lines used in this study.**

**Supplementary Table S13)** lists the genes affected in the RNAseq results from 0-2 hrs old Drosophila embryos mutant for *Top3β*$^{26}$ **and** *FMR1*-. All combinations of elevated and lower reads were considered. Genes with the following expression level changes were selected: adjusted p-value < 0.0002 and absolute log2-fold changes >1.

**Supplementary Table S14)** Selected pairs of transcript isoforms from the same gene that show reciprocal expression levels in *Top3β*$^{Y332F}$ and *Top3β*$^{26}$, but (with one exception) not in not *Top3β*$^{\Delta RGG}$.
